# PAXIP1 and STAG2 converge to maintain 3D genome architecture and facilitate promoter/enhancer contacts to enable stress hormone-dependent transcription

**DOI:** 10.1101/2022.12.27.521987

**Authors:** Isabel Mayayo-Peralta, Sebastian Gregoricchio, Karianne Schuurman, Selçuk Yavuz, Anniek Zaalberg, Aleksander Kojic, Nina Abbott, Bart Geverts, Suzanne Beerthuijzen, Joseph Siefert, Tesa M. Severson, Martijn van Baalen, Liesbeth Hoekman, Cor Lieftink, Maarten Altelaar, Roderick L. Beijersbergen, Adriaan B. Houtsmuller, Stefan Prekovic, Wilbert Zwart

## Abstract

How steroid hormone receptors (SHRs) orchestrate transcriptional activity remains only partly understood. Upon activation, SHRs bind the genome and recruit their co-regulators, crucial to induce gene expression. However, it remains unknown which components of the SHR-recruited co-regulator complex are essential to drive transcription following hormonal stimuli. Through a FACS-based genome-wide CRISPR screen, we comprehensively dissected the Glucocorticoid Receptor (GR) co-regulatory complex involved in gene-target regulation. We describe a novel functional cross-talk between PAXIP1 and the cohesin subunit STAG2 that is critical for regulation of gene expression by GR. Without altering the GR cistrome, PAXIP1 and STAG2 depletion alter the GR transcriptome, by impairing the recruitment of 3D-genome organization proteins to the GR complex. Importantly, we demonstrate that PAXIP1 is required for stability of cohesin on the genome, its localization to GR-occupied sites, and maintenance of enhancer-promoter interactions. Moreover, in lung cancer, where GR acts as tumor suppressor, PAXIP1/STAG2 loss enhances GR-mediated tumor suppressor activity by modifying local chromatin interactions. All together, we introduce PAXIP1 and STAG2 as novel co-regulators of GR, required to maintain 3D-genome architecture and drive the GR transcriptional programme following hormonal stimuli.

## INTRODUCTION

Ligands with a cholesterol-backbone bind with high affinity to different Steroid Hormone Receptors (SHRs), leading to modulation of transcriptional networks related to various cellular functions. Upon activation, SHRs bind the chromatin by engaging with hormone-responsive elements (HREs), predominantly found in enhancers, located distally from promoters^1, 2^. Once localized to their binding sites, SHRs are able to recruit a large complex of co-regulators required for initiation of SHR-dependent transcription^3, 4^. SHRs, together with their co-regulator complex, loop toward their target promoters, establishing enhancer-promoter interactions and subsequently leading to gene regulation^3, 4^. SHRs play pivotal roles in human disease, including cancer. This is best exemplified in breast cancer, prostate cancer, and childhood leukemia where Estrogen Receptor alpha (ER*α*), Androgen Receptor (AR) and Glucocorticoid Receptor (GR), respectively^5–7^ serve as focal points for targeted therapy that successfully reduces disease burden^2, 8, 9^.

SHR-mediated transcription relies on the assembly of a transcriptional complex on HREs. First, pioneer transcription factors such as FOXA1 bind to condensed chromatin, displacing linker histones, demarcating and rendering regulatory elements accessible for SHRs to bind^6, 10^. Once SHRs occupy their cognate regulatory elements, a coordinated recruitment of co-regulators occurs, including chromatin remodeling factors, mediator complex, histone modifiers including histone lysine (de)-methyl transferases (KMT/KMDs), histone acetyltransferases (HATs) and ultimately RNA polymerase II (RNAPII) to drive gene transcription^11–17^. Altogether, SHR activation ultimately results in a large-scale transcriptional complex that loops towards targeting promoters to regulate gene expression, relying on 3D genome architectural proteins, such as the cohesin complex^18–20^. As a result, SHR and their interactors, together with the transcriptional machinery and 3D-genome organization proteins converge to either induce or repress specific target genes^12, 17, 21, 22^.

Although different studies addressed the cross-talk, interaction, and co-localization of SHRs with their co-regulators^12, 17, 21–23^, little is known about which interactors are required for SHR activity, versus those that act redundantly. By using Rapid Immunoprecipitation of Mass spectrometry of Endogenous proteins (RIME)^24^, or quantitative-multiplexed RIME (qPLEX-RIME)^25^ most interactors of different SHRs including ER*α*, AR and GR have been described ^21, 25–31^. However, the latter only provides information of all interactors but not of the functional contribution nor essentiality of every recruited SHR-interactor on the transcriptional output.

In this study, we sought to comprehensively identify which co-regulators are essential to drive SHR-dependent transcription in a gene-specific manner, using GR as a model. We performed a genome-wide CRISPR screen followed by FACS sorting to identify which proteins affect GR-mediated activity. Surprisingly only a small number of hits were identified, suggesting that vast majority of GR interactors act redundantly. We did, however, discover a novel functional cross-talk between PAXIP1 and the cohesin complex, that to our knowledge, has never been described before. PAXIP1, a subunit of the histone modifier complex KMT2D/C^32, 33^, acts in a non-canonical manner to regulate GR-mediated gene expression regulation by functionally interacting with STAG2; a member of the 3D-genome architecture complex cohesin. Finally, we report PAXIP1 to be essential for stability of cohesin on chromatin and its localization to GR-occupied sites, maintenance of enhancer-promoter interactions following hormonal stimuli, and to ensure a fully-functional GR transcriptional program.

## RESULTS

### Genome-wide CRISPR screen identifies PAXIP1 and STAG2 as essential regulators of GR-mediated transcription

Previously, we comprehensively annotated the GR-interacting protein repertoire in various lung cancer cell lines, including A549 cells^31^. However, which of these GR interacting proteins are essential for GR function remains unknown. To identify proteins critical for GR-action in a comprehensive manner, we set up a FACS-based genome-wide CRISPR screen, in which we used the expression of a classical GR target gene – *FKBP5* – as proxy of GR activity (**Fig 1A** and **Fig Sup. 1A-D**). Exposure of the lung cancer cell line A549 to glucocorticoids (GCs) increased GR binding at the *FKBP5* locus **(Fig Sup. 1 A)**, and induced transcription of *FKBP5* over time **(Fig Sup. 1B)**. Importantly, *FKBP5* mRNA and protein levels were not induced upon GC treatment in GR-knockout cells **(Fig Sup. 1C,D)**. Jointly, these controls confirm the GR-dependent status of FKBP5, reinforcing its role as marker gene for GR activity. In this screen, A549 cells were transduced with the CRISPR genome-wide Brunello library, including 77,441 sgRNAs, with an average of four targeting sgRNAs per gene, and 1000 non-targeting controls^34, 35^. Cells were subsequently selected with puromycin, and cultured for 14 days prior to stimulation. In order to induce GR-mediated transcription, A549 cells were treated with GCs for 24 hours, subsequently fixed and stained for protein expression of FKBP5. Cells were FACS-sorted for approximately 7.5% of low (FKBP5^low^) and 7.5% high FKBP5 expression (FKBP5^high^) populations **(Fig. 1A)**. Next, genomic DNA was isolated and sequenced to detect abundance of sgRNAs in both populations. Through MaGECK analyses^36^, we compared the sorted FKBP5^high^ and FKBP5^low^ populations, to identify which genes are individually essential to drive GR activity **(Fig. 1B)**. These analyses identified two critical positive controls – *NR3C1* (encoding for GR) and *FKBP5* – as top hits **(Fig. 1B)**, demonstrating the robustness of our screen. Importantly, we identified *PAXIP1* and *STAG2* among the top-enriched hits in the FKBP5^low^ population **(Fig. 1B)**, implying a critical role of these proteins in regulating GR action. To validate our findings, we generated *PAXIP1* and *STAG2* CRISPR-mediated gene disruption models **(Fig. 1C)**. Loss of either *PAXIP1* or *STAG2* significantly reduced the expression of FKBP5, both at the protein **(Fig. 1D)** and RNA level **(Fig. 1E and Fig Sup. 1E)**. Importantly, GR protein levels remained unaltered in both *PAXIP1*-KO and *STAG2*-KO cells **(Fig. Sup. 1F)**, demonstrating that the observed reduction of FKBP5 expression is not a consequence of altered GR protein stability or transcription.

**Figure 1.**
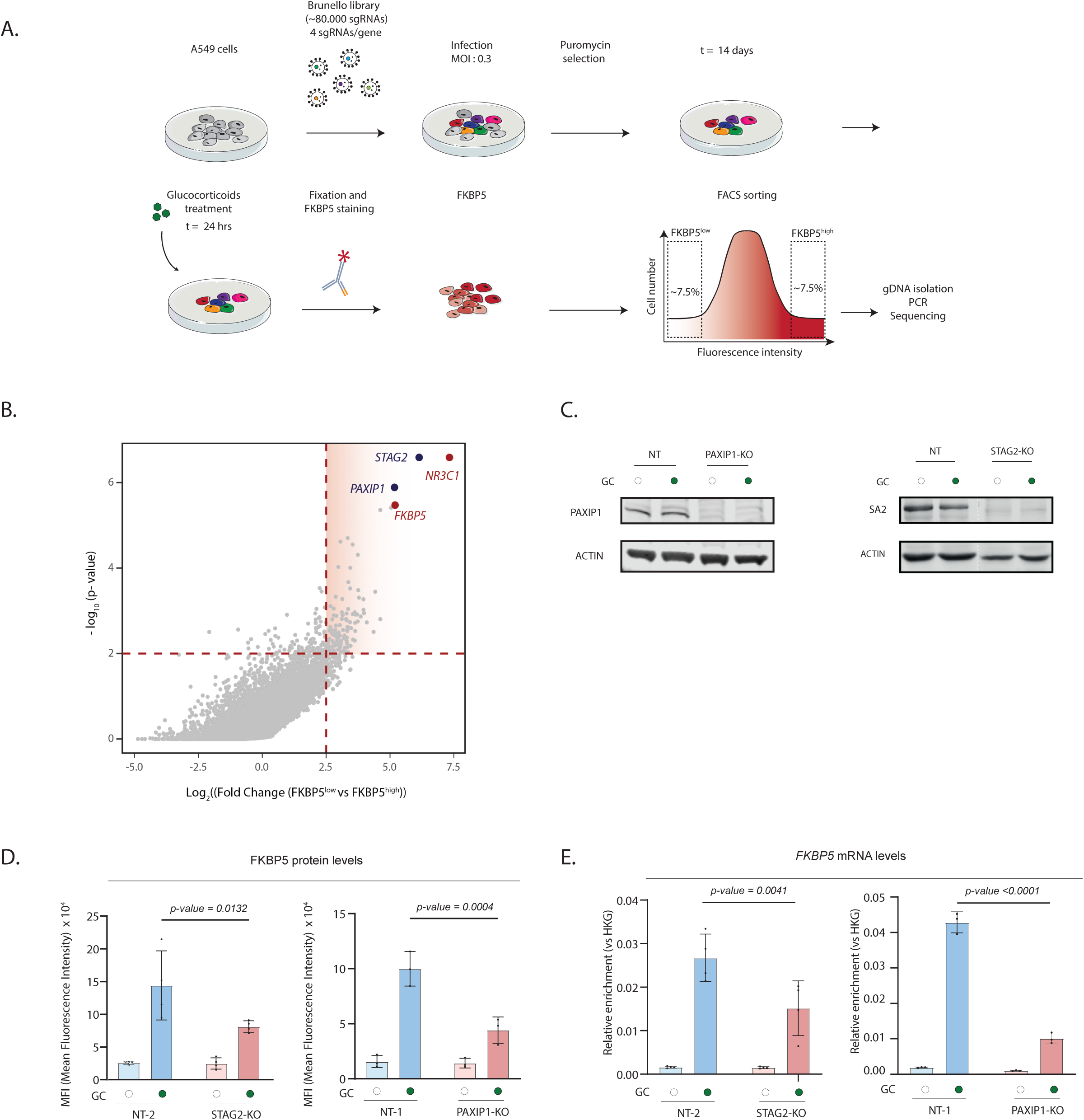
PAXIP1 and STAG2 are crucial for GR-mediated transcription. A. Schematic representation of the FACS-based genome-wide CRISPR screen to identify regulators of GR function. B. Screen results: scatter plot of log_2_ (Fold Change [FKBP5^low^ vs FKBP5^high^ expression]) vs -log_10_(p-value). Dotted lines indicate threshold values (log_2_(Fold Change)>2.5 & - log_10_(p-value)>2). Analyses were performed using MaGECK. Red highlighted dots depict *NR3C1* and *FKBP5*, serving as positive controls. Blue highlighted dots indicate other significant hits of the screen. C. Western blot analyses showing PAXIP1 (left) and SA2 (protein encoded by STAG2, right) expression, with actin as control, in NT, *PAXIP1-KO* and *STAG2*-KO cells (n =2). Cells have been treated with DMSO (empty dot) or GCs (filled dot) for 24 hours. D. FKBP5 protein levels in A549 cells depleted of *PAXIP1* and *STAG2* expression assessed by Flow Cytometry analyses. Quantification of Mean-Fluorescence Intensity (MFI) of FKBP5 expression (n=4 in NT vs *STAG2*-KO, n=3 in NT vs *PAXIP1*-KOs). Cells have been treated with DMSO (empty dot) or GCs (filled dot) for 24 hours. Mean values ± SD are depicted. Two-way ANOVA test was performed. E. Relative *FKBP5* mRNA levels (normalized to geometric mean of housekeeping genes *GAPDH* and *ACTIN*) of DMSO (empty dot) and GC-treated (filled dot) in cells depleted of *PAXIP1* and *STAG2* expression. Mean values ± SD are depicted (n=4 biological replicates in NT vs *STAG2*-KO, n=3 technical replicates in NT vs *PAXIP1*-KOs). Two-way ANOVA test was performed.

### PAXIP1 as a novel functional interactor of STAG2

Having identified PAXIP1 and STAG2 as novel regulators of GR function, we next aimed to elucidate the mechanism behind this observation. For this purpose, we first assessed a possible impact of these proteins on GR/DNA interactions, through chromatin immunoprecipitation followed by sequencing (ChIP-seq) analyses for GR in non-target (NT), *PAXIP1*-KO and *STAG2*-KO cells, both in vehicle and GC-treated conditions. As expected for NT cells, GR chromatin interactions were induced by GC-treatment (**Fig. 2A and 2B**). Interestingly, GR binding to the *FKBP5* locus was not affected in neither *PAXIP1*-KO or *STAG2*-KO cells (**Fig. 2A**), suggesting that PAXIP1 and STAG2 regulate GR functionality downstream of its capacity to associate with the chromatin. The latter was, importantly, also observed on a genome-wide scale (**Fig. 2B**). As expected, genomic distribution analyses showed that GR binding occurs mainly at introns and distal intergenic regions in all cell lines (**Fig. Sup 2A**). However, we did observe a slight enrichment of GR localization at promoters and promoter-proximal regions, relative to most-closely located transcription start site (TSS) in *PAXIP1*-KO cells when compared to NT control (**Fig. Sup. 2B**). In addition, motif analyses at peaks identified in NT, *PAXIP1*-KO and *STAG2*-KO cells showed, as expected, glucocorticoid response elements (NR3C1), JUN and forkhead transcription factors (FOX) among the top most enriched motifs, with no major differences between models (**Fig. Sup. 2C**).

**Figure 2.**
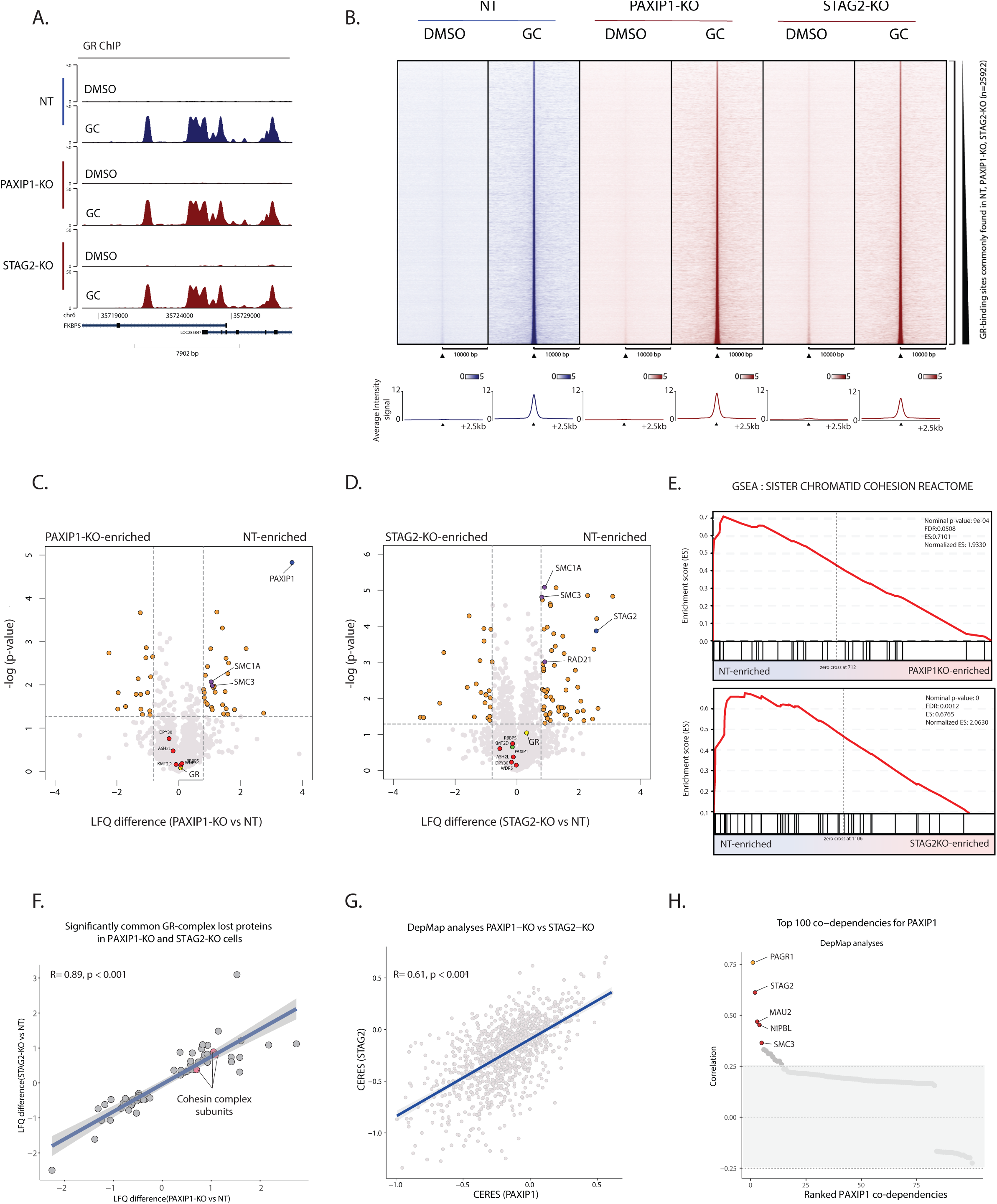
PAXIP1 functionally interacts with Cohesin. A. Snapshot of GR binding around the *FKBP5* locus in non-targeting (NT), *PAXIP1*-KO, and *STAG2*-KO cells, comparing vehicle (DMSO) and treated (GC) conditions. Data represent the average of three biological replicates (n=3). B. Heatmap (top) and average density plot (bottom) of GR ChIP-seq signal in NT, *PAXIP1*-KO, *STAG2*-KO cells in vehicle (DMSO) or treated (GC) conditions. Data are centered at GR-peaks shared in NT, *PAXIP1*-KO and *STAG2*-KO in GC-treated arm, depicting a ±10kb (heatmap) and ±2.5kb (average plot) window around the peak center. Data represent the average of three biological replicates. C. RIME analyses showing the composition of the GR protein complex in *PAXIP1*-KO cells. Volcano plot comparing GR-IP in NT cells over *PAXIP1*-KO. Cells were treated with GCs for 2 hours. Proteins considered to be interacting with GR are highlighted in orange. Significance cut-off is depicted with a dotted line (LFQ>0.8, -log(p-value)>1.3) (n=4). D. RIME analyses depicting the composition of the GR protein complex in *STAG2*-KO cells. Volcano plot comparing GR-IP in NT cells over *STAG2*-KO. Cells were treated with GCs for 2 hours. Proteins considered to be interacting with GR are represented in orange. Significance cut-off is represented with a dotted line (LFQ>0.8 -log(p-value)>1.3) (n=4). E. GSEA enrichment profiles for Sister chromatid cohesion reactome (M27181), comparing NT vs *PAXIP1*-KO (top) or NT vs *STAG2*-KO (bottom). Ranking is based on the LFQ values of GR-RIME data. Nominal p-value and NES were determined with GSEA. F. Correlation plot between those proteins that were commonly significantly (-log(p-value)>1.3) differentially recruited to the GR complex (Pearson correlation R=0.89, p-value <0.001). G. Correlation between the DepMap CERES values of *STAG2* and *PAXIP1* across all 1043 cell lines from the DepMap database (Pearson correlation R= 0.61, p<0.001). H. DepMap analyses showing top 100 co-dependencies for *PAXIP1*. Pearson correlation values are depicted.

Due to the critical role of these co-regulators on GR transcriptional activity, we next investigated whether the composition of the GR complex was altered upon PAXIP1 or STAG2 depletion. To do so, we performed Rapid Immunoprecipitation of Endogenous proteins (RIME) mass spectrometry analyses^24^ for GR in both PAXIP1 (**Fig. 2C**) and STAG2 (**Fig. 2D**) deficient cells. PAXIP1 is a subunit of the multiprotein complex KMT2D/C; a histone methyl transferase complex, responsible for methylation of H3K4, demarcating enhancers^32, 33, 37^. Surprisingly, GR-mediated recruitment of the KMT2D/C complex (DPY30, ASH2L, KMT2D, WDR5, RBBP5) was not affected when PAXIP1 was knocked out (**Fig. 2C**). In line with this, depletion of a main subunit of the complex – KMT2D – did not affect expression of FKBP5, as measured by flow cytometry analyses (**Fig. Sup. 2D**). Jointly, these findings suggest that PAXIP1 impacts GR action in a KMT2D/C-independent manner in lung cancer cells.

Surprisingly, GR RIME analyses showed diminished recruitment of members of the 3D-genome organization-related cohesin complex in cells lacking PAXIP1, including SMC1A and SMC3 (**Fig. 2C**). These proteins serve to facilitate promoter/enhancer interactions and enable long-range gene regulation by *cis*-acting transcription factors^37^. These results suggest that PAXIP1 may regulate association of cohesin to GR-bound regulatory sites on the genome. Analogous to our PAXIP1 findings, knockout of STAG2 perturbed the capacity of GR to interact with SMC1A, SMC3 and RAD21 (**Fig. 2D**). The strong similarities of *PAXIP1*-KO and *STAG2*-KO impacting the GR protein interactome were further highlighted in Gene Set Enrichment Analyses (GSEA), in which the proteins group in the sister chromatid reactome dataset were significantly lost upon either *PAXIP1* or *STAG2* knockouts (**Fig. 2E** and **Fig Sup 2E**). Reduced recruitment of cohesin subunits to the GR complex upon depletion of either *PAXIP1* or STAG2, suggested a possible novel functional crosstalk of PAXIP1 with STAG2. These findings were further confirmed in quantitative analyses on our RIME data, demonstrating that core GR-associated proteins remained recruited even upon *PAXIP1* and *STAG2* ablation (**Fig Sup 2F**). Furthermore, high correlation was found in the levels of differential protein recruitment to the GR complex in NT when compare to *STAG2*-KO and *PAXIP1*-KO cell lines, including cohesin subunits (**Fig. 2F**). To gain further insights into a possible PAXIP1-STAG2 functional interaction, we analyzed the essentiality of *STAG2* and *PAXIP1* in a publicly available CRISPR-screen dataset containing data for 1,043 cell lines from different tumor types and derived from various organs (DepMap 21Q4 public; https://depmap.org/portal/) (**Fig. 2G**). Interestingly, we found a high level of co-dependency between PAXIP1 and STAG2 across all cell lines (R = 0.61, p-value <0.001) (**Fig. 2G**), implying that PAXIP1 and STAG2 might be involved in regulating similar cellular processes. Next to STAG2, cohesin loaders MAU2 and NIPBL^38^, as well as the subunit of cohesin SMC3, were all found amongst the highest co-dependencies for PAXIP1 in this dataset (**Fig. 2H**), providing further evidence for a functional crosstalk between PAXIP1 and STAG2. All together, we find that loss of both PAXIP1 and STAG2 does not affect GR chromatin binding, but does diminish the recruitment of proteins involved in 3D-genome organization to the GR complex, which is associated with a loss of GR target-gene regulation, as measured by expression of GR-activity marker FKBP5.

### PAXIP1 is required for cohesin stability on the genome, its localization to GR-bound enhancers and enhancer-promoter interactions

Our findings indicated that PAXIP1 may regulate cohesin recruitment to the chromatin-bound GR complex. To further explore this, we endogenously tagged cohesin subunit SMC1 with an EGFP-tag in both wild-type and PAXIP1 deficient cells **(Fig. Sup. 3A)**, and performed recovery after photobleaching (FRAP) experiments to determine SMC1 mobility in live cells **(Fig. 3A and 3B; Fig. Sup. 3B)**. *PAXIP1* knockout increased SMC1-EGFP mobility in these cells, both under vehicle and GC treatment **(Fig. 3A and 3B)**, indicating that cohesin chromatin stability was impaired upon PAXIP1 loss **(Fig. 3A and 3B).** We performed computer simulations of the FRAP curves^39^ and calculated the different immobile and mobile fractions of SMC1-EGFP **(Fig. Sup. 3B)**. Upon knocking out *PAXIP1*, the immobile fraction of cohesin was decreased in both DMSO and GC-treated cells, indicating reduced stability of cohesin on the genome **(Fig. Sup. 3B)**. In addition, when cells were exposed to GCs, the immobile fraction in PAXIP1-KOs was also lower than in NTs, with short and intermediate immobile fractions being more prominent in *PAXIP1*-KOs when compared to NT **(Fig. Sup. 3B)**. All together, we observed that depletion of PAXIP1 results in a general reduction of cohesin stability on the chromatin, both in DMSO and GC-treated cells.

**Figure 3.**
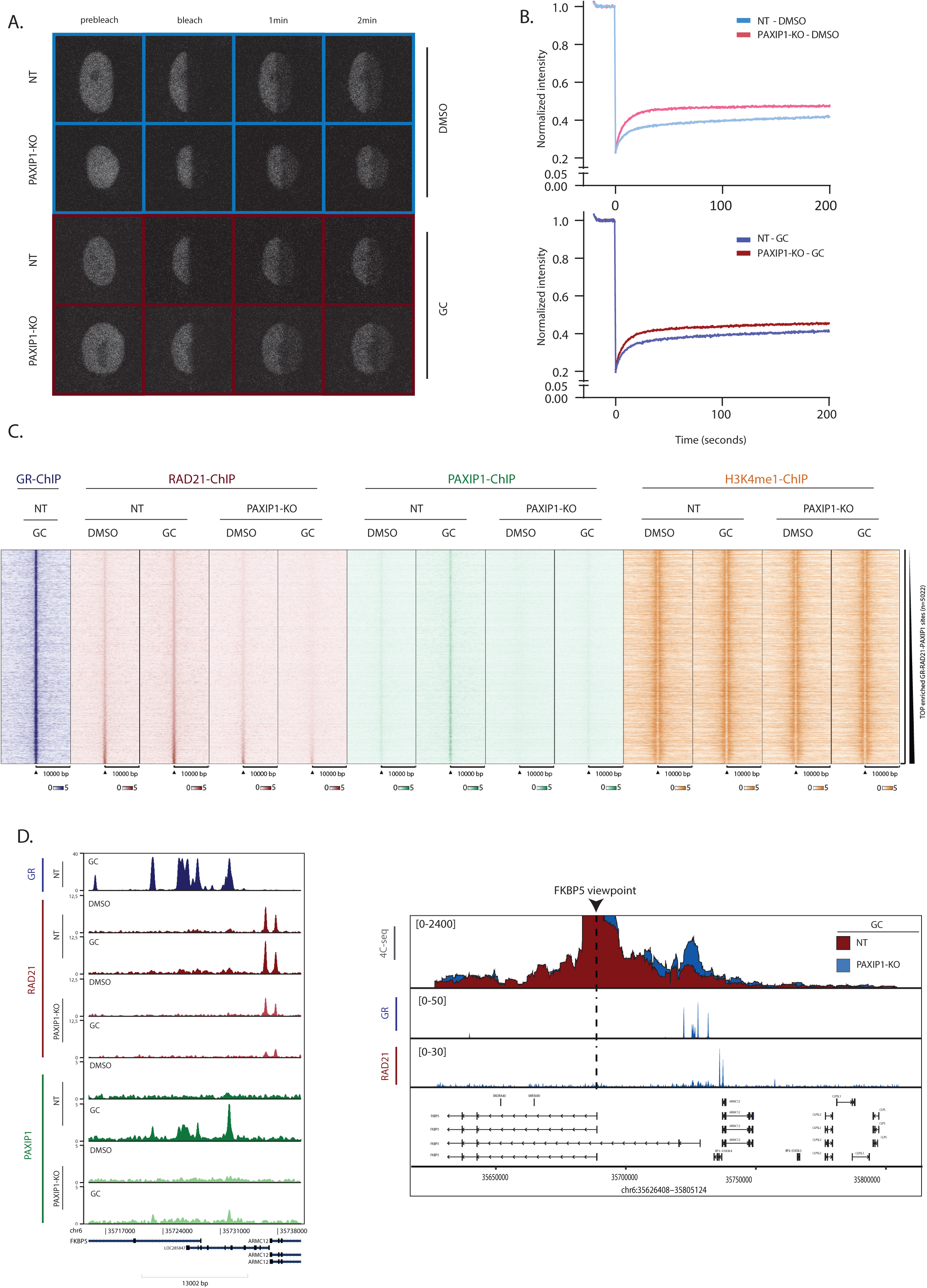
PAXIP1 is required for Cohesin function. A. Representative image of FRAP experiments in NT-SMC1-EGFP and *PAXIP1*-KO-SMC1-EGFP cells with vehicle (top, blue) and GC-treated (bottom, red) cells. B. Quantification of FRAP experiments over time in NT-SMC1-EGFP and *PAXIP1*-KO-SMC1-EGFP cells with vehicle (above) and GC-treated cells (below). Average ± SD for 30 cells quantified over 3 independent biological experiments. Intensity is pre-bleached normalized. C. Heatmap of RAD21, PAXIP1 and H3K4me1 ChIP-seq signal in NT, *PAXIP1*-KO cells in vehicle (DMSO) or treated (GC) conditions. Data are centered at commonly bound RAD21 and GR-peaks in NT cells treated with GC, depicting a ±10kb window around the peak center (n=2). D. Snapshots of GR, RAD21 and PAXIP1 ChIP-seq tracks around the *FKBP5* locus. E. 4C-seq experiments in NT and *PAXIP1*-KO cells using the *FKBP5* TSS as a viewpoint. Cells were treated with GCs for 2 hours. 4C data is shown as the mean of three independent biological experiments. ChIP tracks of RAD21 and GR in GC-treated cells are depicted. ChIP-tracks are a representative snapshot of one biological replicate.

To confirm the FRAP results, we employed an orthogonal approach assessing PAXIP1 and the cohesin subunit RAD21 genome interactions in both wild-type and PAXIP1-depleted cells, by ChIP-seq analyses **(Fig. 3C and Fig. Sup. 3C)**. Both PAXIP1 and RAD21 were found to occupy GR-binding sites, which was dependent on GC-treatment, suggesting both PAXIP1 and RAD21 are recruited to these sites following GR chromatin binding **(Fig. 3C and Fig. Sup. 3C)**. Importantly, RAD21 recruitment to GR-bound sites was lost when PAXIP1 was knocked out **(Fig. 3C,D and Fig. Sup. 3C)**, confirming a critical role of PAXIP1 for RAD21 binding to the genome. Genomic distribution analyses showed that GR-PAXIP1-RAD21 co-occupancy on the genome occurs mainly at introns and distal intergenic regions **(Fig. Sup. 3D**). Motif analyses at GR-PAXIP1-RAD21 sites showed, as expected, glucocorticoid response elements, JUN and forkhead transcription factors (FOX) among the top most enriched motifs **(Fig. Sup. 3E**). Interestingly, when interrogating all RAD21 binding sites across the genome in GC-treated cells, we observed reduced promoter signal for RAD21 upon PAXIP1 knockout **(Fig. Sup. 3F)**, implicating that PAXIP1 enables recruitment of RAD21 to the TSS of genes. In addition, we performed motif analyses on these sites, which were enriched for CTCF, but also for classical steroid hormone receptor motifs (**Fig. Sup. 3G**). In parallel, we also performed ChIP-seq experiments for H3K4me1, which is a histone modification marking for enhancers, generally deposited by KMT2D/C complex^37, 40^ (**Fig. 3C and Fig. Sup. 3H**). Interestingly, enrichment for H3K4me1 at these GR-bound sites was not affected by *PAXIP1* knockout at the *FKBP5* locus (**Fig. Sup. 3H**) and on a genome-wide scale (**Fig. 3C**). This is in line with the unaltered KMT2D complex recruitment to the GR complex in *PAXIP1*-KO cells (**Fig. 2C**). Jointly, these data suggests that cohesin recruitment to GR-binding sites is dependent on non-canonical and novel function of PAXIP1.

As we demonstrated that PAXIP1 is required to localize and stabilize cohesin on the genome, we next sought to investigate whether PAXIP1 is necessary for STAG2 role in maintaining local enhancer-promoter interactions. For this purpose, we performed 4C-seq experiments in cells expressing or lacking PAXIP1 expression, selecting the TSS of *FKBP5* gene as view-point (**Fig. 3E**). In wild-type cells, we found a significant increase of interactions between the *FKBP5* promoter and GR-bound enhancer regions upon stimulation with GCs (**Fig. 3E** and **Fig. Sup. 3I**), which is in line with our previous observations reporting GR-stimulated promoter-enhancer interactions^31^. The interacting regions spanned the genomic coordinates in which we observed co-binding of PAXIP1, RAD21 and GR (**Fig. 3E** and **Fig. Sup. 3I**). However, in cells depleted for PAXIP1, promoter/enhancer interactions were not increased, remaining at basal levels despite GC treatment (**Fig. 3E** and **Fig. Sup. 3I**). Cumulatively, these data illustrate a novel role of PAXIP1 in facilitating stable cohesin-chromatin dynamics, cohesin localization to GR-binding sites and GC-induced enhancer-promoter interactions.

### PAXIP1 deficiency mimics the transcriptional program of STAG2 depleted cells

Since PAXIP1 and STAG2 appear to have a functional cross-talk that is reflected at the level of 3D genome organization, we next examined whether PAXIP1 and STAG2 also control similar transcriptional programs upon GR activation. For this, we generated RNA sequencing (RNA-seq) data in *PAXIP1*-KO and *STAG2*-KO cells, cultured in vehicle or GC-treated conditions (**Fig. 4A**). Both PAXIP1 and STAG2 knockouts affected baseline expression of various genes, suggesting a broader significance of both these factors for gene expression regulation (**Fig. 4A**). Importantly, we observed that GC-mediated gene expression was altered in *PAXIP1*-KO and *STAG2*-KO cells (**Fig. 4A**), implying the importance of PAXIP1 and STAG2 in regulating only a subset of GR-target genes. Importantly, we observed a high correlation (R=0.86, p<0.001) on the level of directionality and magnitude of affected genes following knockout of *PAXIP1* and *STAG2* (**Fig. 4B**), with a highly significant (p-value < 2.2e-16) overlap in the transcriptional changes in GC-treatment conditions (**Fig. Sup. 4A**). As STAG2 is crucial for enhancer-promoter interactions^41^, we interrogated whether the similarities on PAXIP1- and STAG2-dependent transcriptional programs were due to regulation of chromatin looping by PAXIP1 on a genome-wide scale. To test this hypothesis *in silico*, we used sevenC analyses, which allows for the prediction of enhancer-promoter interactions based on CTCF sites and proximal RAD21 binding^42^. Using RAD21 ChIP-seq data in NT and *PAXIP1*-KO cells, we observed the number of predicted loops in GC-treated cells were significantly decreased in *PAXIP1*-KO cells (**Fig. 4C** and **Fig. 4D**; **Fig. Sup 4B**). Next, we aimed to validate these predictions experimentally, and confirm that the changes in gene expression were due to perturbed enhancer-promoter interactions. For this purpose, we selected a GR-target gene – *P4HA3* – whose expression was significantly reduced upon *PAXIP1*-KO in GC-treated cells (**Fig. Sup. 4B** and **4C**). 4C-seq analyses were performed, using the *P4HA3* promoter as a viewpoint and confirmed a loss of enhancer-promoter interactions in the area spanning RAD21 binding, which also coincides with the genomic locus of *PPME1* gene (**Fig. Sup. 4D**). Interestingly, GR-mediated expression of genes in close proximity of the *P4HA3* locus – such as *PPME1* – was also significantly reduced (**Fig. Sup. 4C**), which could suggest regional transcriptomic effects when enhancer-promoter interactions are lost. All together, we demonstrate that PAXIP1 and STAG2 control a similar transcriptional landscape by maintaining enhancer-promoter interactions.

**Figure 4.**
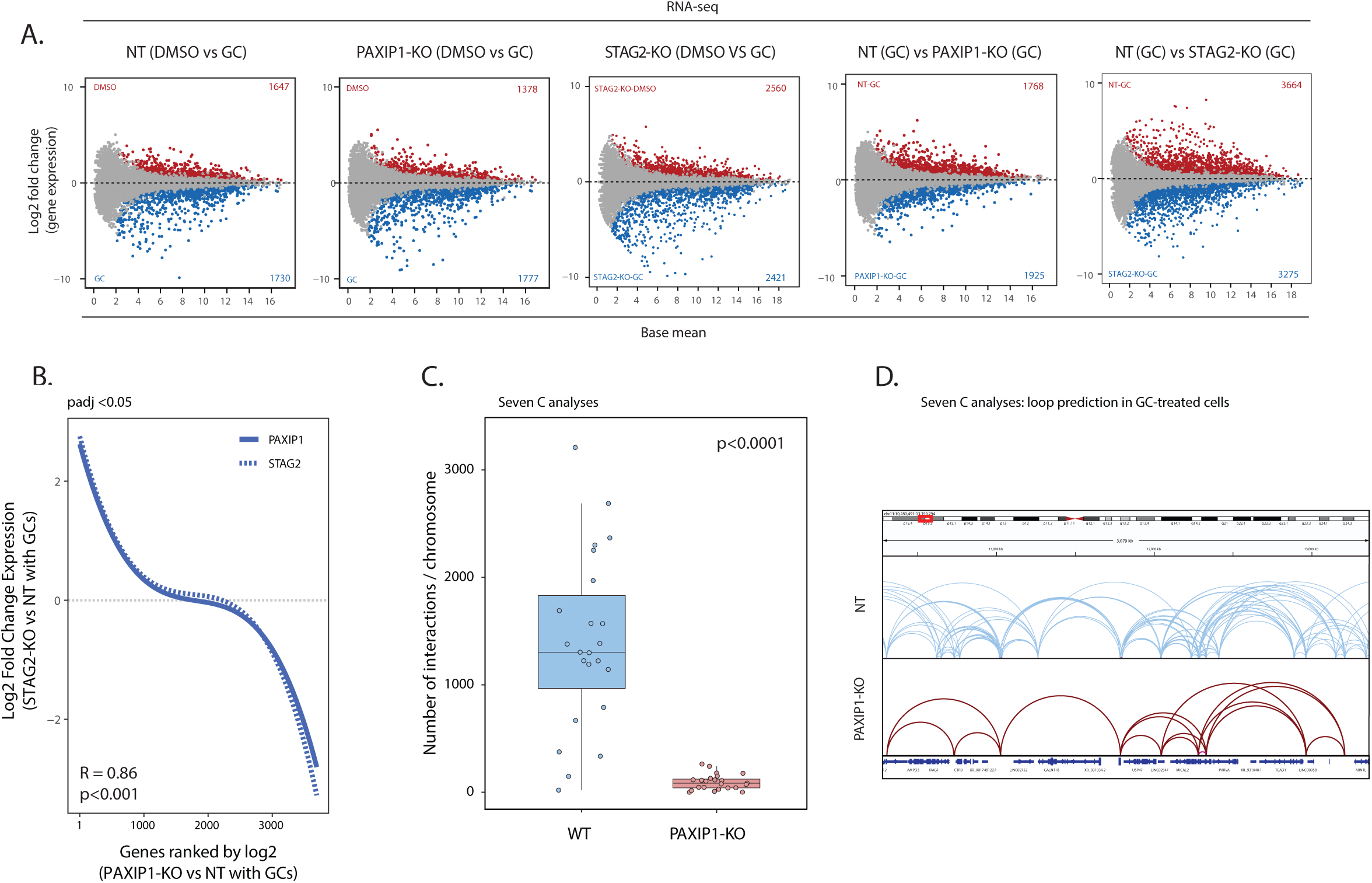
PAXIP1 and STAG2 control similar transcriptional programs. A. MA plots depicting differential gene expression between DMSO- and GC-treatment in NT, *PAXIP1*-KO and *STAG2*-KO cells, NT vs *PAXIP1*-KO in GC-treated cells, and NT vs *STAG2*-KO in GC-treated cells. Red and blue dots indicate significantly differentially expressed genes with log_2_(fold change)>1 and adjusted p-value <0.05. Data represent the average of three independent biological replicates. B. Correlation of differentially expressed genes in *STAG2*-KO cells with *PAXIP1*-KO cells, both in GC-treated conditions. Genes differentially expressed in *PAXIP1*-KO cells (padj < 0.05) were ranked based on log_2_(fold change). C. sevenC analyses of WT and *PAXIP1*-KO based on RAD21 ChIP-seq and CTCF motifs. Number of predicted genomic interactions per chromosome in GC-treated conditions are depicted. One sample t-test was used. P-value < 0.0001 D. Snapshot of sevenC predicted chromosomal interactions in chr 11, both in NT and *PAXIP1*-KO cells.

### Loss of PAXIP1 enhances the tumor suppressor action of GR by modifying local chromatin interactions

In this study, we used the lung cancer model A549 to study GR co-regulators. Importantly, we^31^ and others^8, 43–45^ have previously shown that GC treatment of lung cancer cells reduces tumor cell proliferation, for which we recently reported the mechanisms underlying GR-mediated cancer cell dormancy ^31^. As we observed major changes in the transcriptional landscape of PAXIP1- and STAG2-deficient cells, we next interrogated whether this would result in alterations of tumor cell growth following GR activation. For these purposes, we performed cell proliferation analyses in NT, *PAXIP1*-KO and *STAG2*-KO cells, treated with GCs. In line with our previous findings, GC-treatment diminished tumor cell proliferation capacity (**Fig. 5A** and **Fig. Sup. 5A**). Interestingly, knockout of *PAXIP1* and *STAG2* further strengthened the anti-proliferative effects of GC treatment, while not affecting the cell growth capacity in absence of GCs (**Fig. 5A** and **Fig. Sup. 5A**). These data suggest an enhanced GR-mediated tumor suppressive capacity upon loss of PAXIP1 or STAG2. Based on these results, we hypothesized that loss of PAXIP1 or STAG2 leads to the activation of newly acquired GR-responsive tumor suppressor genes, dampening cell cycle progression upon PAXIP1 and STAG2 depletion. To test this hypothesis, we selectively analyzed genes with significantly gained expression in GC-treated *PAXIP1*-KO and *STAG2*-KO cells, and tested these for overlap with a Cell Cycle geneset (hsa04110), resulting in the identification of two candidate driver genes: *CDKN1B* and *CDKN2C* (**Fig. 5B**). Interestingly, both *CDKN1B* and *CDKN2C* are known to mediate cell cycle arrest^46, 47^. As PAXIP1 depletion resulted in slight increased GR chromatin binding around the *CDKN1B* locus, in contrast to *CDKN2C* (**Fig. Sup. 5B**), we selectively focused follow-up experiments on *CDKN1B* (**Fig. Sup. 5B**). Importantly, *PAXIP1* knock-out resulted in increased *CDKN1B* expression, both on mRNA (**Fig. 5C**), and protein level with increased P27 – encoded by *CDKN1B* – expression (**Fig. 5D**). To confirm that GR-mediated expression of *CDKN1B* was responsible for the enhanced sensitivity to GCs in *PAXIP1*-KO cells, we generated *PAXIP1*/*CDKN1B* double knock-out (DKO) cells (**Fig. Sup. 5C**). We observed that *PAXIP1*/*CDKN1B*-DKO cells restored the tumor cell proliferative phenotype that was lost upon *PAXIP1* knock-out, rendering the cells again less responsive to GCs (**Fig. 5E**).

**Figure 5.**
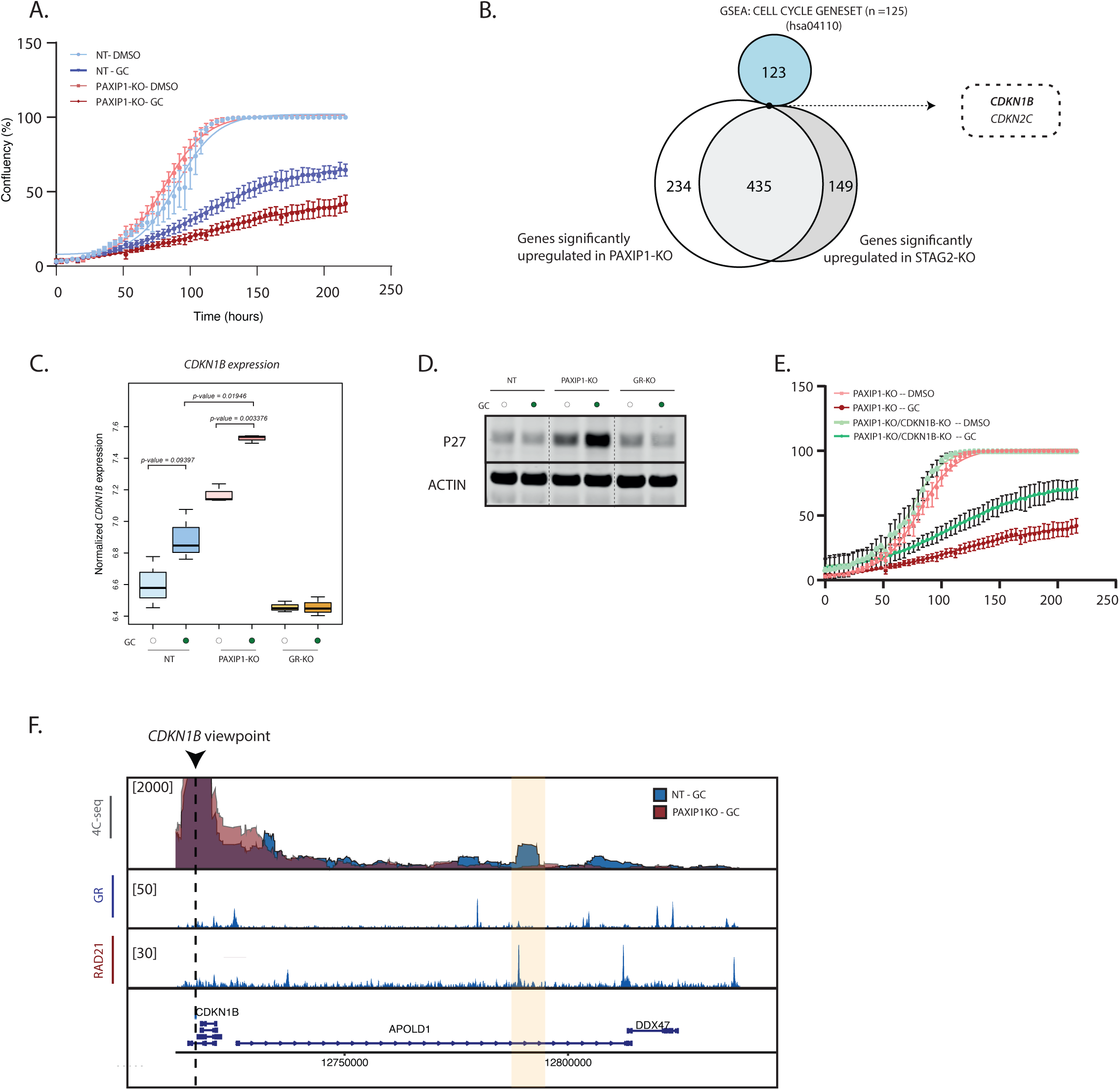
Loss of STAG2 and PAXIP1 enhance the tumor suppressor action of GR by modifying local chromatin interactions. A. Cell proliferation of NT and *PAXIP1*-KO cells following DMSO or GC treatment over time (n = 3 up to 168 hrs, n = 2 from 168hrs to 216hrs, error bars represent 95% confidence interval) B. Overlap of significantly differentially expressed genes in *PAXIP1*-KO and *STAG2*-KO cells following GC stimulation (log_2_(FoldChange) > 0.75; padj < 0.05), with a cell cycle gene set (hsa04110). C. Boxplot showing normalized *CDKN1B* expression in NT, *PAXIP1*-KO and *GR*-KO cells upon vehicle or GC treatment for 8 hours (n =3). p-value was calculated using U Mann-Whitney t-test. D. P27 protein expression levels in NT, *PAXIP1*-KO and *GR*-KO cells upon vehicle or GC treatment for 24 hours. Actin was used as a loading control (n = 2). E. Cell proliferation of *PAXIP1*-KO cells and *PAXIP1*/*CDKN1B*-DKO following DMSO or GC treatment over time (n = 3 up to 168hrs, n = 2 from 168hrs to 216hrs, error bars represent 95% confidence interval). F. 4C-seq experiments in NT and *PAXIP1*-KO cells using the *CDKN1B* TSS as a viewpoint showing NT vs *PAXIP1*-KO in GC-treated conditions. Cells were treated for 2 hours with GCs. 4C data is shown as the mean of three independent biological experiments. ChIP tracks of RAD21 and GR in GC-treated cells are depicted. ChIP-tracks are a representative snapshot of one biological replicate.

As GR binding to the promoter of *CDKN1B* was not altered in *PAXIP1*-KO or *STAG2*-KO cells (**Fig Sup 5B**), we next investigated impact of *PAXIP1* knockout on 3D genome regulation around the *CDKN1B* locus. For that, we performed 4C-seq experiments using the *CDKN1B* promoter as viewpoint and observed again alterations upon PAXIP1 loss, but now this is associated with increased target gene expression (**Fig. 5F**). A loss of long-range chromatin interactions was observed stemming from the *CDK1NB* promoter, spanning a region containing both GR and RAD21 binding sites. These data suggest that this loop might act as an insulator between the promoter and the GR binding site, allowing GR-mediated transcription of *CDKN1B* upon PAXIP1 loss, resulting in loop disruption.

Cumulatively, we observed that loss of PAXIP1 and STAG2 enhances the response of lung cancer cells to GCs, mediated by alterations in long-range 3D-genome contacts and resulting gain of a novel target gene.

## DISCUSSION

SHRs are ligand-sensing transcription factors that play critical roles in a large variety of cellular processes, including maintenance of cellular homeostasis, metabolism, immune system activity, sex dimorphism and cell proliferation^8^. SHRs are expressed throughout the body, often in a highly tissue-selective manner, and their deregulation can result in different human pathologies such as metabolic disorders and cancer development^48^. SHRs regulate gene expression through specific DNA regions, generally enhancers, along with a large complex of co-regulators^3^. Although it is known that SHRs can interact with a large number of proteins, little is known about which ones are essential for their activity.

In this study, we use GR function in lung cancer as a model system, and interrogate on a genome-wide level which co-regulator proteins are required for GR activity. Due to the design of our screen, we exclusively identify hits that are not replaceable in their ability to shape GR-mediated gene expression, suggesting that many of the classical GR interacting proteins may be functionally redundant, and could be compensated by other components in the complex. Interestingly, we identified both PAXIP1 and STAG2 as two key essential proteins in facilitating GR-mediated gene expression, that converge to control 3D genome architecture. We demonstrate that PAXIP1 is required for enhancer-promoter interactions mediated by STAG2, necessary to drive GR-mediated gene expression.

We propose a mode of the functional crosstalk between PAXIP1 and STAG2 that is as follows: upon activation, GR and PAXIP1 bind together to regulatory elements, inducing enhancer-promoter interactions, for which cohesin is necessary (**Fig. 6A and 6B**). We identified PAXIP1 as essential for recruitment of cohesin to GR-binding sites and consequently, to maintain enhancer-promoter interactions. Upon loss of PAXIP1, there is a reduction of localization of cohesin to GR binding sites and this results in impaired GR-mediated transcription (**Fig. 6A**). On the other hand, we propose that GR can also bind together with PAXIP1 and cohesin to GR-binding sites resulting in an insulator loop, avoiding enhancer/promoter contacts to prevent GR-mediated transcription, or prevent basal levels of gene expression (**Fig. 6B**). However, in the absence of PAXIP1, cohesin localization to those sites is lost, resulting in a loss of insulator loop formation and consequently gained gene expression (**Fig. 6B**).

**Figure 6.**
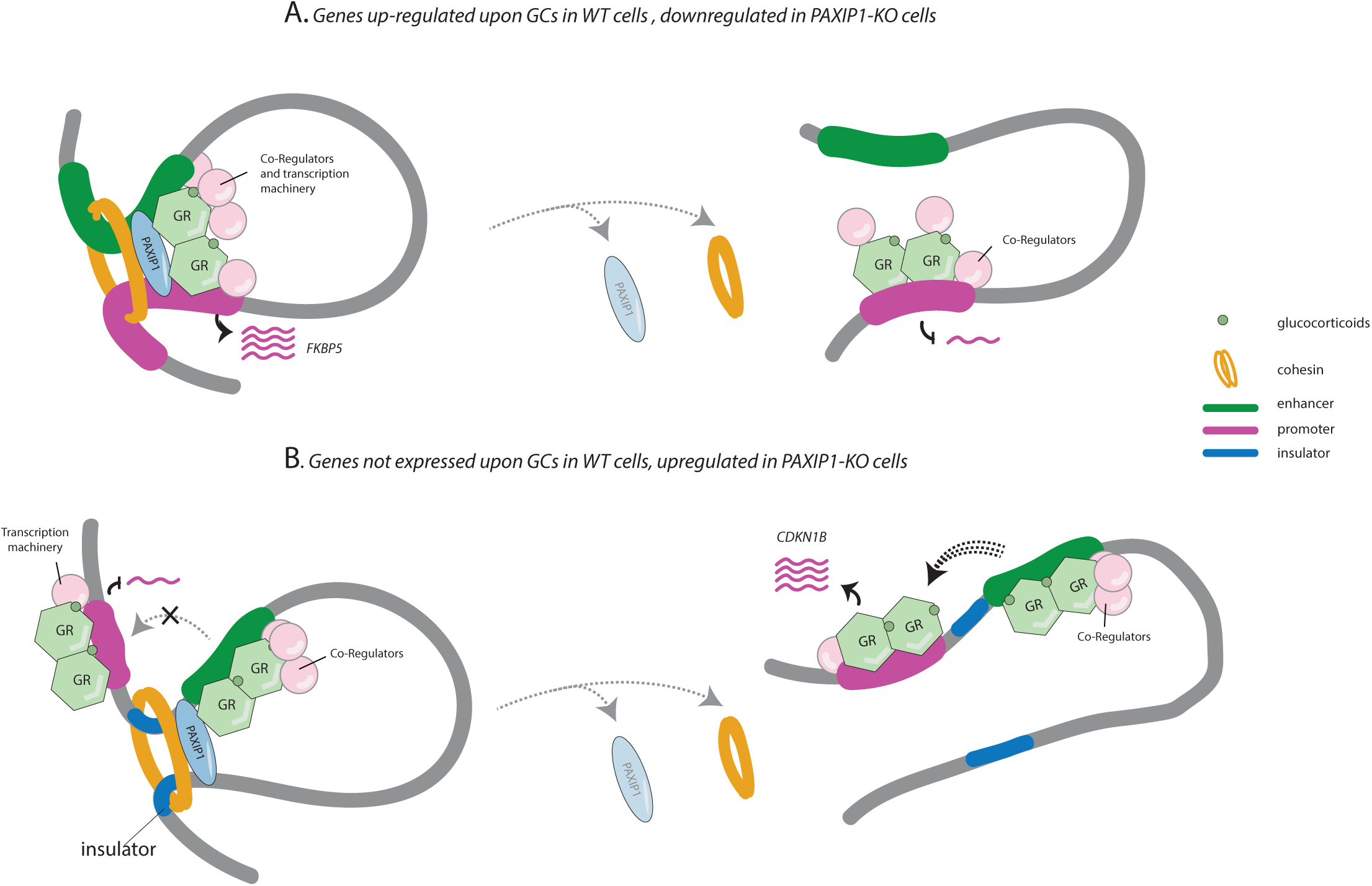
Proposed model. PAXIP1 and STAG2 converge to facilitate enhancer-promoter interactions and fine-tune GR-target gene expression either by enhancing expression of GR-target genes (A) or by blocking expression through insulator loops (B)

STAG2 belongs to the ring-shaped structured cohesin complex, which plays an essential role in sister chromatid cohesion, DNA damage repair, as well as the formation and stability of *cis*-chromatin loops for transcriptional regulation^49–51^. Here, we report a new role of PAXIP1 regulating cohesin function. We demonstrate that PAXIP1 is required for cohesin stability on chromatin and its localization to GR-bound sites. While the functional role of PAXIP1 in 3D genome organization is novel, prior studies have reported an interaction between 3D-genome organization proteins and different SHRs, including GR and ER*α*, being able to bind specific genomic regions together with cohesin^20, 52–55^. Moreover, GR activation increases both the number and intensity of enhancer-promoter interactions of their target genes upon hormonal stimulation^20^. GR was recently described to interact with NIBL, which is responsible for cohesin loading onto DNA, localization of cohesin to GR-bound sites, and promotion of long-range genomic interactions^18^.

Here we show, that cohesin binding to GR-bound sites is critically dependent on PAXIP1, with its loss altering GR-mediated transcriptome. Importantly, PAXIP1 and STAG2 depletion resulted in similar changes in gene expression, providing further evidence for their functional interplay. PAXIP1 is a component of the KMT2D/C complex^32, 33^, a histone methyltransferase necessary for H3K4 methylation at enhancers of actively transcribed genes^40, 56^. Surprisingly, we observed that in PAX1P1-depleted cells, GR-mediated recruitment of the KMT2D complex and its subunits was not altered. Moreover, H3K4me1 levels remained unaffected upon PAXIP1 knockout, and depletion of KMT2D did not alter expression of the GR-target protein FKBP5. Altogether, this suggests that PAXIP1 can function in a KMT2D-independent manner to regulate cohesin binding and stability on the genome and subsequently regulate enhancer-promoter interactions following hormonal-response. In line with this, prior reports describe PAXIP1 to act in a KMT2D/C-independent manner in conjunction with its counterpart PAGR1 ^57^. PAXIP1 has been involved in immunoglobin class switching and V(D)J recombination^58, 59^ ; a biological process in which cohesin-mediated loop extrusion plays a key role^60, 61^. Moreover, PAXIP1 is critically involved in DNA damage response, localizing together with 53BP1 in DNA damage sites, to enable ATM-mediated phosphorylation of cohesin subunit SMC1 and ensuring a correct DNA damage response^62, 63^. Finally, in parallel to our study, Van Schie et al, described the interplay of PAXIP1 and PAGR1 to maintain chromatin-bound cohesin during cell cycle progression^64^. These observations not only support our findings of PAXIP1 regulating cohesin activity in a KMT2D/C-independent manner, but also highlight a more-general contribution of PAXIP1 function on cohesion action, beyond its role on facilitating promoter/enhancer interactions, and re-positions PAXIP1 as a critical component of cohesin action.

Although our study is focused on the functional crosstalk between PAXIP1 and STAG2 following GR activation in lung cancer, we hypothesize that our results may be extrapolated to other tissues and *cis*-acting transcription factors, which is supported by the more-general action of cohesin in facilitating transcription factor action in 3D genomic space, beyond GR alone. We observed a high level of co-dependency between PAXIP1 and the cohesin subunits STAG2, SMC1A and cohesin loaders MAU2 and NIBPL across >1000 cell lines, suggesting that the PAXIP1-cohesin crosstalk is underexplored, yet serves a universal synergistic role across many tissues and tumor types. Therefore, interrogating whether the activity of other *cis*-acting transcription factors – including NF-*κ*B or other SHRs – are dependent on PAXIP1-cohesin functional crosstalk would be of high interest for future studies.

GR is known to function as a tumor suppressor in lung cancer ^31^. Here we show that depletion of PAXIP1 or STAG2 results in enhanced tumor suppressor activity of GR, due to increased *CDKN1B* expression. The latter encodes for P27; a well described tumor suppressor, which is often downregulated in cancer and serves as a prognostic marker in lung cancer ^47^. We describe a possible mechanism, by which changes in enhancer-promoter interactions proximal to the *CDKN1B* locus, result in acquired regulation of *CDKN1B*. We hypothesize, that this altered *CDKN1B* expression a consequence of disruption of an insulator loop following PAXIP1 and STAG2 depletion, which warrants further investigation.

Regulation of gene expression in 3D genomic space is a research field in active development, in which the cohesion complex – originally reported to drive mitotic entry and orchestrate chromosomal segregation – plays a pivotal role. Now, we reveal that PAXIP1 – originally reported as part of the KMT2D/C complex – functionally and critically contributes to cohesin action, driving expression of genes under control of a *cis*-acting transcription factor, the Glucocorticoid Receptor.

## ACKNOWLEDGEMENTS

We thank members of the Zwart and Bergman labs for valuable feedback, suggestions and input. This work was supported by KWF/Alpe d’Huzes and Oncode Institute. We would like to thank the NKI genomics core facility for next-generation sequencing and bioinformatics support. We thank the NKI Flow Cytometry facility for technical support. We thank the NKI Proteomics/Mass Spectrometry facility (M.A. and L.H. are supported by the Dutch NWO X-omics Initiative). We thank Reuven Agami, Julien Champagne and Remco Nagel for fruitful discussions and technical support.

## AUTHOR CONTRIBUTIONS

Conceptualization: I.M.P., S.P., and W.Z.; S.P. and W.Z. were responsible for project funding. I.M.P., S.G., and A.K. performed 4C-seq experiments and S.G. analyzed 4C-seq experiments; I.M.P., K.S., S.P., N.A. and S.B. performed RIME, ChIP-seq and RNA-seq experiments; I.M.P. and N.A. performed RT-qPCR and western blot analyses; I.M.P., K.S., A.Z., M.v.B performed the CRISPR screen; J.S. and T.M.S. provided bioinformatics support for peakcalling ; L.H. and performed and analyzed the proteomic experiments. ; C.L. and R.L.B. provided bioinformatics and technical support for the CRISPR screen; S.Y., B.G. and A.B.H. performed FRAP experiments. I.M.P., S.P. and W.Z. wrote the manuscript, with input from all co-authors.

## DECLARATION OF INTERESTS

The authors declare no competing interests

## METHODS

### Cell lines

Lung cancer cell line A549 was obtained from American Type Culture Collection (ATCC). A549 cells were cultured in Dulbecco’s Modified Eagle Medium (DMEM)/F12 (1:1) (1x) + Glutamax (Life technologies) supplemented with 10% fetal calf serum (FCS), 1% penicillin-streptomycin (Pen/Strep) (5000 U/mL, life technologies), unless otherwise stated. All cell lines were cultured at 5% CO_2_ and 37 °C. All cell lines were genotyped and tested negative for mycoplasma.

### FACS-based genome-wide CRISPR screen

The human CRISPR Brunello library^34, 35^ was used in this study, which was a kind gift from Roderick Beijersbergen (The Netherlands Cancer Institute, NKI). The Brunello CRISPR lentiviral library was transduced into A549 cells at a low multiplicity of transduction (MOI) of ∼0.3 to ensure that only one sgRNA was incorporated per cell. Transduced cells were subsequently selected with 2μg/mL of puromycin for 3 days and left untreated for 14 days. After that, cells were incubated in DMEM/F12 (1:1) (1X) + Glutamax, supplemented with dextran-coated charcoal-treated FCS (DCC) for 24 hours and then treated with 1μg/mL of hydrocortisone (HC) for additional 24 hours. After treatment, cells were fixed in 70% ethanol and left overnight at 4 °C. Fixed cells were washed with PBS and permeabilized with PBS containing 0.25% triton X-100 (Sigma-Aldrich) and left on ice for 15 min. Cells were stained with a recombinant anti-FKBP5 antibody (AlexaFluor 647, ab198979, 1:400) in 1% bovine serum albumin (BSA)-PBS. Stained cells were washed with PBS and resuspended in 1%BSA-PBS. Cells were then FACS sorted for 7.5% FKBP5^high^ or 7.5% FKBP5^low^ of the population, with ∼3.500.000 cells/arm to ensure a 500x coverage of the library. Genomic DNA was subsequently isolated using Gentra Puragene Cell kit according to manufacturer’s instructions. gRNAs were amplified by two consecutive PCRs as previously described (Korkmaz et al., 2016). DNA libraries were sequenced on HiSeq 2500 platform (single-read; 65bp). Read counts were normalized, relative total sizefactors were calculated and that the values within a sample were divided by the respective sample sizefactor, reads of each replicate were pooled together and subsequent analyses were performed using MaGeCK, using default parameters (v0.5.9.4^36^). Putative GR regulators were identified comparing the sgRNA abundance amongst the 7.5% FKBP5^high^ vs 7.5% FKBP5^low^ populations and a robust rank aggregation (RRA) score was determined for each gene. Those genes with an FDR<0.15, log_2_(fold change)>2.5 and –log_10_(p-value)>2 were considered as putative hits. Results of the screen can be found on Supplementary Table 1. FACS-sorting gating strategy can be found in Extended Data 1.

### Genome editing

#### CRISPR knockout cell lines

For the generation of CRISPR knockout cell lines, guide RNAs were selected from the GeCKOv2 human CRISPR KO library to target *PAXIP1* (CACCGGTGATTCTGTCCGTTCAGTG) *STAG2* (AG-TCCCACATGCTATCCACA), non-targeting (NT-1) (CACCGAACTACAAGTAAAAGTATCG), non-targeting 2 (NT-2) (GTATTACTGATATTGGTGGG), *NR3C1* (GTGAGTTGTGGTAACGTTGC) and *CDKN1B* (GGGTTAGCGGAGCAATGCGC). The selected gRNAs were cloned into the lentiCRISPRv2 plasmid as previously described^66^. All constructs were verified by Sanger sequencing. Lentivirus was produced by transfection of viral packaging vectors and Lentiv2-CRISPR constructs in HEK293T cells. On the first day, 3.5 million HEK293T cells were plated in 10 cm^2^ dishes and incubated overnight. On day two, viral vectors were produced by mixing three packaging constructs in a 1:1:1 ratio: pRC/CMV-rev 1B, pHDM-G, and pHDM-Hgpm2, after which 10.5 µg of the packaging mix was added to 17.5 µg of every single CRISPR construct together 105 µL of polyethylenimine (PEI, 1 mg/mL), incubated for 15 min and added to HEK293T cells. After an overnight incubation, cells were refreshed with 8 mL of medium. Next day, the supernatant was harvested and added onto A549 cells together with polybrene (200x, 1.6g/L). After 48hrs, cells were selected with either puromycin (2μg/mL) or blasticidin (1mg/mL) SMC1-EGFP tagged cell lines In order to generate endogenously expressing EGFP-tagged SMC1A cell lines, we first selected the most suitable 20bp-long sequence close to the C-terminal end of the SMC1A gene for gRNA binding. The gRNA sequences were designed by using an online algorithm (http://crispor.tefor.net) and the gRNA closest to the SMC1’s stop-codon were chosen (FW: AAAATACTGCTACTGCTCAT; RV: ATGAGCAGTAGCAGTATTTT). The selected gRNAs were ordered as single-stranded DNA oligos (IDT) and annealed according to an earlier described protocol^67^. A PX459 vector (pSpCas9-2A-Puro V2.0; Addgene 62988)^68^ was selected for inducing DSBs via CRISPR/Cas9. To do this, the PX459 vector was first digested by BbslI restriction enzyme (NEB: R0539S) and afterwards the oligos were ligated by T4 DNA ligase (NEB: M0202S). Plasmids were sequenced by Sanger sequencing to confirm the cloning of the gRNA sequences.

In parallel, gene fragments were synthesized (IDT) containing 470 bp long homology arms flanked on both sides of the gRNA target site and excluding SMC1’s stop-codon. Subsequently, a donor template with the EGFP-stop-codon cassette including a 6 times glycine-alanine spacer sequence was cloned in between the two homology arms by using MluI (NEB: R0198S) and BglII (Roche: 10567639001) restriction enzymes. Fragments were ligated and afterwards inserted into a pCR-Blunt II TOPO backbone vector (Thermofisher: K280002). Donor template plasmid sequences were confirmed by Sanger sequencing. Next, PX459 and donor template were transfected using lipofectamine 3000 (Thermofisher) according to manufactureŕs instructions. Next, cells were selected with puromycin (2ug/mL), clones were selected and genotyped.

### Flow Cytometry

Cells were cultured in DMEM/F12 (1:1) (1X) + Glutamax, supplemented with dextran-coated charcoal-treated FCS (DCC) with DMSO or 1mg/mL hydrocortisone (HY-N0583, MedChemExpress) for 24 hours. Cells were fixed with 70% ethanol at 4°C overnight. Cells were washed with PBS, permeabilised with 0.25% Triton X-100 in PBS-T for 15 minutes on ice, and stained with primary recombinant Alexa Fluor® Anti-FKBP51 antibody (ab198979, Abcam, 1:100) diluted in 1% BSA-PBS or left unstained. Cells were washed and resuspended in 1%BSA/PBS solution. Cells were sorted using Attune TM NxT Acoustic Focusing Cytometer (Thermo Fisher Scientific). Single-cell flow cytometry analysis was performed using FlowJo™ Software version 10.7.1 (BD Biosciences)

### RNA isolation, reverse-transcription and quantitative real-time PCR (RT-qPCR)

Cells were cultured in DMEM/F12 (1:1) (1X) + Glutamax, supplemented with dextran-coated charcoal-treated FCS (DCC) with DMSO or 1mg/mL hydrocortisone (HY-N0583, Med-ChemExpress) for 24hrs. Total RNA was isolated using Invitrogen™ TRIzol™ Reagent (15596026, Thermo Fisher Scientific) according to manufactureŕs instructions. First-strand cDNA was synthesized from 1 µg of isolated RNA by using SuperScript^®^ III First-Strand Synthesis System for Reverse Transcriptase-PCR (18080-051, Life Technologies). RT-qPCR was performed using SensiMix™ SYBR® No-ROX Kit (QT650-05, Bioline) in a QuantStudio™ 6 Flex System (Thermo Fisher Scientific), and analyzed using the QuantStudio Software. Primers can be found in Supplementary table 2.

### Western blot

Cells were lysed using 2x Laemmli buffer (120 mM Tris, 20% glycerol, 4% SDS) supplemented with protease inhibitor (1:100) and phenylmethylsulfonyl fluoride (PMSF, 1:200) upon over-night treatment with 100nM dexamethasone (HY-14648, MedChemExpress) or 1mg/mL hydrocortisone (HY-N0583, MedChemExpress) or left untreated. Lysates were sonicated (EpiS-hear Probe Sonicatore, Active Motif) for 10 cycles with one second intervals and a 20% amplitude. Equal amounts of protein per lysate were run for one hour at 100 V on an 8% acrylamide gel (MilliQ, 40% acrylamide, 1.5 M Tris pH 6.8, 10% SDS, 10% APS, TEMED) in SDS-PAGE 1x Running buffer (25 mM Tris, 0.25 M glycine, 0.1% SDS). Proteins were transferred on ice at 100 V for 90 min or at 0.9 mA overnight at 4 °C on a nitrocellulose membrane in cold 1x Transfer buffer (24 mM Tris, 192 mM glycine). Membranes were stained with Ponceau S (Thermo Fisher) and subsequently blocked in 3% BSA (A8022, Sigma/Merck) in 1x PBS-Tween (137 mM NaCl, 10 mM Na_2_HPO4, 1.5 mM KH_2_PO_4_, 2.6 mM KCl, 0.1% Tween-20) for one hour and incubated with primary antibodies against GR (12041, Cell Signaling Technology, 1:1000), PAXIP1 (ABE1877, Merck, 1:1000), STAG2 (A300-158A, Bethyl, 1:1000), Actin (MAB1501R, Merck, 1:1000), HSP90 (sc-13119, Santa Cruz Biotechnologies, 1:1000), P27 (610242,BD Biosciences, 1:1000), SMC1 (Bethyl:A300-055A; 1:1000) diluted in 3% BSA/PBS-T for two hours. After three washing steps in PBS-T, membranes were incubated with secondary antibodies donkey-α-mouse 680 RD (926-68073, LI-COR Biosciences, 1:10.000), donkey-α-rabbit 800 CW (926-32213, LI-COR Biosciences, 1:10.000) and donkey-α-goat 680 RD (926-68074, LI-COR Biosciences, 1:10.000), diluted in 3% BSA/PBS-T for one hour. Membranes were scanned and analysed using an Odyssey^®^ CLx Imaging System (LI-COR Biosciences) and ImageStudio™ Lite v.5.2.5 software (LI-COR Biosciences).

### Rapid immunoprecipitation of endogenous proteins (RIME)

RIME was performed as previously described^24^. In brief, cells were treated for two hours with 1mg/mL hydrocortisone. Subsequently, cells were fixed and crosslinked with 1% formaldehyde (15714, Electron Microscopy Sciences) for exactly 10 minutes, quenched with glycine (0.125 M) and washed three times with PBS. Cells were collected in 1xPBS supplemented with 1X complete EDTA-free protease inhibitor cocktail (PI tablets, 5056489001, Roche) on ice. Cells were lysed as previously described (Mohammed et al., 2016), and cell lysates were sonicated for six cycles (30 sec on/30 sec off) using the Bioruptor^®^ Pico (B01060001, Diagenode). Per RIME, 50 µL of magnetic Protein A beads (10008D, Thermo Fisher Scientific) beads were conjugated with either 15 µL anti-GR (12041, Cell Signaling Technology) rotating overnight at 4 °C.

For mass spectrometry, peptide mixtures were prepared and measured as previously described^27^, with the following exceptions. For GR RIMEs in NT vs *PAXIP1*-KO cells, peptide mixtures (10% of total digest) were loaded directly onto the analytical column and analyzed by nanoLC-MS/MS on an Orbitrap Fusion Tribrid mass spectrometer equipped with a Proxeon nLC1200 system (Thermo Scientific). Solvent A was 0.1% formic acid/water and solvent B was 0.1% formic acid/80% acetonitrile. Peptides were eluted from the analytical column at a constant flow of 250 nl/min in a 120-min gradient, containing a 104-min stepped increase from 6% to 32% solvent B, followed by a 16-min wash at 90% solvent B. For GR RIMEs in NT vs STAG2-KO cells, peptide mixtures (10% of total digest) were loaded directly onto the analytical column and analyzed by nanoLC-MS/MS on an Orbitrap Exploris 480 Mass Spectrometer equipped with a Proxeon nLC1200 system (Thermo Scientific). Solvent A was 0.1% formic acid/water and solvent B was 0.1% formic acid/80% acetonitrile. Peptides were eluted from the analytical column at a constant flow of 250 nl/min in a 90-min gradient, containing a 74-min stepped increase from 6% to 32% solvent B, followed by a 16-min wash at 90% solvent B.

Raw data were analyzed by MaxQuant (GR RIMEs in NT vs PAXIP1-KO cells: version 2.0.1.0; GR RIMEs in NT vs STAG2-KO cells:version 1.6.17.0)^69^ using standard settings for label-free quantitation (LFQ). MS/MS data were searched against the Swissprot Human database (GR RIMEs in NT vs PAXIP1-KO cells: 20,395 entries, release 2021_04; GR RIMEs in NT vs STAG2-KO cells: 20,379 entries, release 2021_01) complemented with a list of common contaminants and concatenated with the reversed version of all sequences. The maximum allowed mass tolerance was 4.5ppm in the main search and 0.5Da for fragment ion masses. False discovery rates for peptide and protein identification were set to 1%. Trypsin/P was chosen as cleavage specificity allowing two missed cleavages. Carbamidomethylation was set as a fixed modification, while oxidation and deamidation were used as variable modifications. LFQ intensities were Log2-transformed in Perseus (GR RIMEs in NT vs PAXIP1-KO cells: version 1.6.15.0; GR RIMEs in NT vs STAG2-KO cells: version 1.6.14.0), after which proteins were filtered for at least 3 out of 4 valid values in at least one sample group. Missing values were replaced by imputation based on a normal distribution (width: 0.3 and downshift: 1.8). Differentially expressed proteins were determined using a Student’s *t*-test.

### Chromatin immunoprecipitation followed by sequencing (ChIP-seq)

Chromatin immunoprecipitation followed by sequencing (ChIP)-seq was performed as previously described^70^. Cells were treated with 100nM Dexamethasone (HY-14648, MedChemExpress) for 2 hours. Cells were fixed in 1% formaldehyde (1039991000, Merck) for 10 min and quenched with 0.125M glycine. Nuclear lysates were extracted as previously described^70^ and sonicated for 13 cycles (30 sec on/30 sec off) using the Bioruptor^®^ Pico (B01060001, Diagenode). The antibodies that were used are the following: GR (12041, Cell Signaling Technology), RAD21 (05-908, Merck), PAXIP1 (ab70434, abcam), H3K4me1 (ab8895, abcam). Per ChIP, 50 µL of magnetic Protein A beads (10008D, Thermo Fisher Scientific) beads were conjugated to 7.5µL of GR antibody, or 5µg of RAD21, PAXIP1, and H3K4me1 antibodies. Immunoprecipitated DNA was processed for library preparation (KAPA library preparation kit, KK8234, Roche). In the case of GR and RAD21 ChIPs, generated libraries were sequenced on the Illumina HiSeq2500 platform (single-end, 65bp reads). For PAXIP1 and H3K4me1 ChIPs, samples were sequenced on Illumina Novaseq 6000 (paired-end, 51bp). For those samples processed in Novaseq, adapter trimming was performed before alignment using seqpurge^71^. All reads were aligned to the Human Reference Genome (GRCh38.102) using Burrows-Wheeler Aligner (v0.7.17; ^72^) and mem algorithm. In samples processed with Novaseq, duplicates were marked umi-aware, using rumidup (https://github.com/NKI-GCF/rumidup). For samples processed in HiSeq, data did not have an UMI, and based in coordinates, biological duplicates were removed. Reads were filtered based on MAPQ quality ≥20 (samtools v1.9), and duplicate reads were removed. Peak calling over input was generated by using MACS2 (v2.1.2; ^73^), by using the peakcalling pipeline https://github.com/csijcs/snakepipes . Consensus peaks between biological replicates were generated using mspc tool^74^.For visualization, mapped reads of replicate samples were merged using SAMtools (v1.10;^75^). Genome browser snapshots, average density plots and tornado plots were generated using Easeq (v1.101; ^76^). Genomic distribution analyses were performed using ChIPseeker (v1.26.2; ^77^) under R 4.0.3. Motif enrichment analyses were performed using the SeqPos motif tool on Galaxy Cistrome^78^.

### Cell proliferation analyses

Cells were plated in a 384-well plate at a density of 250 cells/well. Cells were treated with 100nM of Dexamethasone. Cells were imaged every 4 hours by using an IncuCyte ZOOM Live-Cell Analysis System, and cell confluency percentage was calculated using the incucyteZoom software.

### Circularized Chromosome Conformation Capture sequencing (4C-seq)

Cells were treated with 100nM Dexamethasone (HY-14648, MedChemExpress) for 2 hours. Experiments of 4C-seq were performed as previously described ^79^. Briefly, for each restriction enzyme (RE) combination (RE1 NlaIII, RE2 DpnII; New England Biolabs), replicate and treatment about 10×10^6^ cells were collected, pelleted and cross-linked by 2% methanol-free formaldehyde for 10 minutes. Nuclei were isolated and permeabilized to allow digestion of the chromatin by the primary RE (NlaIII, New England Biolabs). Chromatin fragments were then diluted and ligated before crosslinking reversion. Purified DNA was digested by the secondary RE (DpnII, New England Biolabs) and circularized again by ligation. Re-purified circular fragments were amplified by PCR with View-Point specific primers (see Supplementary Table 2) using the Expand Long Template PCR System (Roche). Resulting amplicons were purified with a 0.8× ratio of AMPure XP beads (Beckman Coulter) and amplified using standard indexed Illumina primers as previously described^79^ using the Expand Long Template PCR System (Roche).

Second-round PCR products were purified with PCR purification columns (Qiagen) and quantified by 2100 Bioanalyzer (Agilent, DNA 7500 kit). 4C library was prepared by equimolar mix of the samples. The pooled library was cleaned-up by AMPure XP beads (0.8× ratio) to remove PCR dimers before sequencing through Illumina MiSeq with a 75bp Single-End reads setup. Fastq files have been demultiplexed by *Cutadapt* (http://journal.embnet.org/index.php/emb-netjournal/article/view/200) and mapped on Hg38/GRCm38 genome assembly and signal normalized by *pipe4C* (v1.1) R-package ^79^ in “*cis*” mode and with default parameters. Score mean of the replicates and plots of normalized mean 4C-seq and ChIP-seq signal were generated in an R v4.0.3 environment by using *get.single.base.score.bw* and *genomic.track* functions from *Rseb* (v0.3.0) (https://github.com/sebastian-gregoricchio/Rseb)(^80^) package in combination with *ggplot2* (v3.3.5) and *ggforce* (v0.3.3) packages.

### Fluorescence recovery after photobleacing (FRAP)

FRAP was performed on a Leica TCS SP8 microscope equipped with a 63x/1.40 NA HC PL APO CS2 oil immersion objective in combination with a 50mW Argon excitation laser using the 488nm line. FRAP measurements were conducted on approximately 50% of the nuclear area (344 pixel by 344 pixel, 100nm pixel size, 2 times line averaging). After 40 scans (500ms interval), a high intensity laser beam at 488nm was utilized for 3 iterations with a 197ms interval to photobleach all GFP locally inside the selected ROI. Subsequently, the bleached ROI was continuously scanned for 1200 iterations with a 500ms interval. Quantitative FRAP analysis was performed using a Monte Carlo simulation environment for modeling complex biological molecular interaction networks^39^.

Computer modelling used to generate FRAP curves for fitting was based on Monte Carlo simulation of diffusion and binding to immobile elements (representing chromatin binding) in an ellipsoidal volume (representing the nucleus). Bleaching simulation was based on experimentally derived three-dimensional laser intensity profiles, which determined the probability for each molecule to become bleached considering their 3D position relative to the laser beam. Diffusion was simulated at each new time step *t* + Δ*t* by deriving a new position (*x_t_*_+Δ*t*_, *y_t_*_+Δ*t*_, *z_t_*_+Δ*t*_) for all mobile molecules from their current position (*x_t_*, *y_t_*, *z_t_*) by *x_t_*_+Δ*t*_=*x_t_ +* G(*r_1_*), *y_t_*_+Δ*t*_=*y_t_ +* G(*r_2_*), and *z_t_*_+Δ*t*_=*z_t_ +* G(*r_3_*), where *r_i_* is a random number (0 ≤ *r_i_* ≤ 1) chosen from a uniform distribution, and G(*r_i_*) is an inversed cumulative Gaussian distribution with μ=0 and σ^2^=2*D*Δ*t*, where *D* is the diffusion coefficient. Immobilization was derived from simple binding kinetics: *k_on_*/*k_off_* = *F_imm_ /* (1–*F_imm_*), where *F_imm_* is the fraction of immobile molecules. The probability per unit time to be released from the immobile state was given by *P_mobilise_* = *k_off_* =1 / *T_imm_*, where *T_imm_* is the characteristic time spent in immobile complexes expressed in unit time steps. The probability per unit time for each mobile particle to become immobilized (representing chromatin-binding) was defined as *P_immobilise_* = *k_on_* = (*k_off_* · *F_imm_*) */* (1– *F_imm_*), where *k_off_* = 1 / *T_imm_*. Note that *k_on_* and *k_off_* in this model are effective rate constants with dimension s^−1^.

In all simulations, the size of the ellipsoid was based on the size of the measured nuclei, and the region used in the measurements determined the size of the simulated bleach region. The laser intensity profile using the simulation of the bleaching step was derived from confocal images stacks of chemically fixed nuclei containing GFP that were exposed to a stationary laser beam at various intensities and varying exposure times. The unit time step Δ*t* corresponded to the experimental sample rate of 100 milliseconds.

For quantitative analysis of the FRAP data, raw FRAP curves were normalized to pre-bleach values and the best fitting curves (by ordinary least squares) were selected from a large set of computer simulated FRAP curves in which three parameters representing mobility properties were varied: diffusion rate, immobile fraction and time spent in immobile state. Because individual curves generated by Monte Carlo modelling, in contrast to analytically derived curves, show the slight variation typical for diffusion of a limited number of molecules in a small volume, we did not use the best-fitting curve only, but took the ten best-fitting curves and calculated the average diffusion coefficients and rate constants corresponding to these curves.

### Genetic dependency analyses

The dependency data used in this study was obtained from DepMap 21Q4 public (https://depmap.org/portal/). Dependency data for *PAXIP1* and *STAG2* genes across 1046 cell lines was downloaded. Pearson correlation coefficient was calculated between CERES scores of PAXIP1 and STAG2. Moreover, top100 co-dependencies for PAXIP1 were downloaded from https://depmap.org/portal/. All data was plotted using ggplot2 (v.3.3.5) and ggpubr (v.0.4.0).

### sevenC analyses

Prediction of loops was performed using sevenC (v1.19.0)^42^, run with default parameters. We used CTCF sites identified using JASPAR 2022 database in the hg38 human genome (“AH104716”), filtered for those with a p-value < 1e-6. As ChIP input, we used peak called files from the RAD21 ChIP performed in NT and *PAXIP1*-KO cells in GC-treated conditions (See ChIP section for more details). One sample t-test was used for analyses.

### RNA-seq

Cells were treated with Dexamethasone (100nM) for 8 hours. Total RNA was extracted using RNeasy Mini kit (Qiagen, Germany) following manufactureŕs instructions. The quality and quantity of the total RNA were assessed by the 2100 Bioanalyzer using a Nanochip (Agilent, USA). NT-1, PAXIP1-KO and GR-KO RNA-seq series was sequenced in the Illumina HiSeq2500 platform. NT-2, STAG2-KO series was sequenced in Illumina Novaseq 6000. Sequencing data were aligned to the human reference genome hg38 using HISAT2 (v2.1.0;^81^), and the number of reads per gene was measured with HTSeq count (v0.5.3;^82^). Read counting, normalization and differential gene expression were performed using R package DESeq2 (v.1.30.1 ;^83^)

---

**Figure Sup 1.**
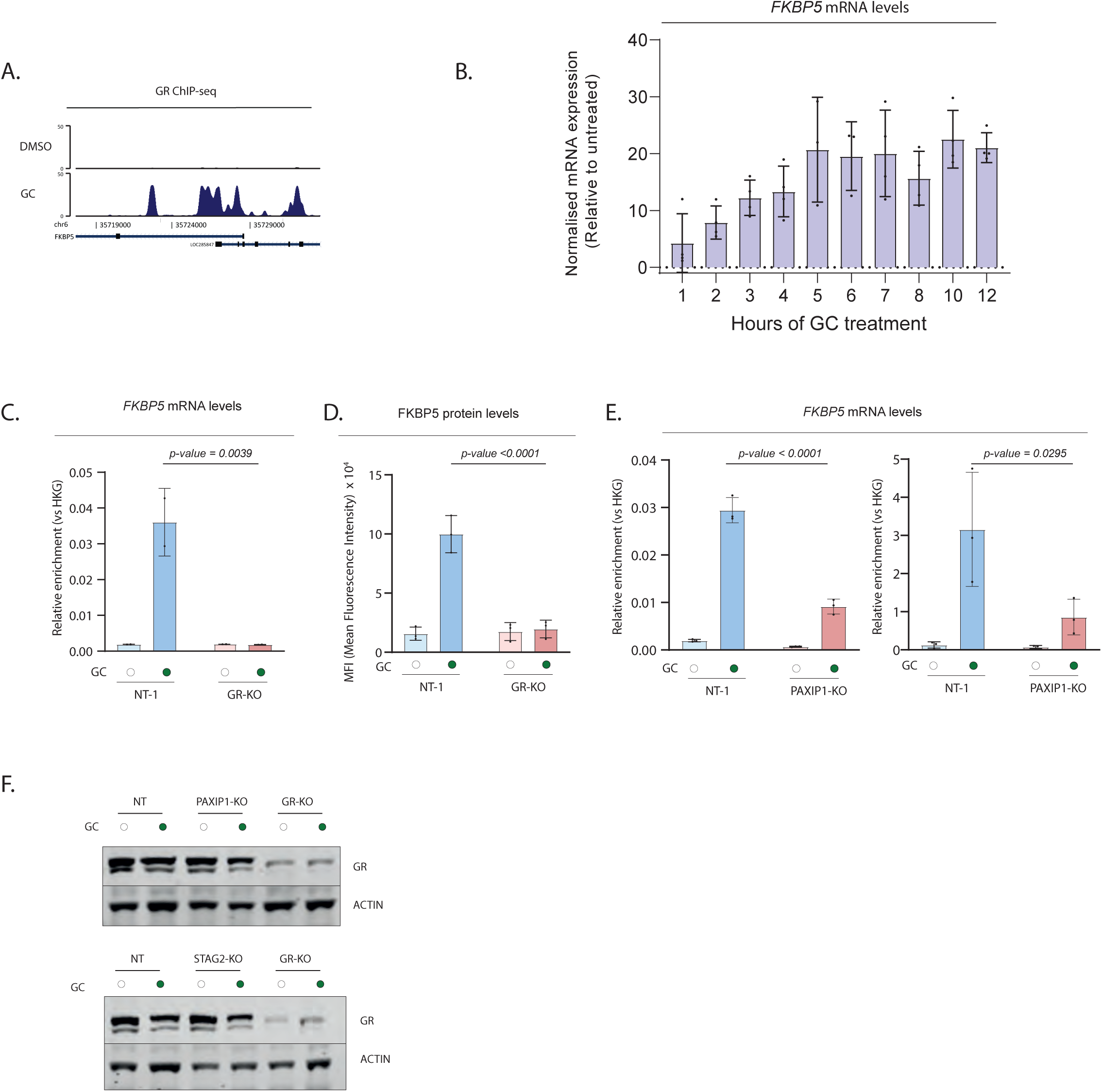
FKBP5 as a proxy of GR-function. Validation of CRISPR screen. A. ChIP-seq track at *FKBP5* locus depicting GR binding following GC treatment for 2 hours. Data is the merge of three independent biological replicates. B. Normalized *FKBP5* mRNA expression levels during a time course experiment of A549 cells exposed to GC treatment (ENCSR897XFT). Mean values ± SD are depicted. (n = 4, for all time points expect t=5,6 hours which is n=3). C. Relative *FKBP5* mRNA levels (normalized to geometric mean of housekeeping genes: *GAPDH* and *ACTIN*) of vehicle (and GC-treated) cells in cells depleted of GR expression. Mean values ± SD are depicted (n=3). D. FKBP5 protein levels in A549 cells depleted of GR expression assessed by Flow Cytometry analyses. Empty and filled dots indicate treatment with DMSO or GCs for 24 hours, respectively. Quantification of Mean-Fluorescence Intensity (MFI) of FKBP5 expression (n=3). Two-way ANOVA test was performed. E. Relative *FKBP5* mRNA levels (normalized to geometric mean of housekeeping genes *GAPDH* and *ACTIN*) of DMSO (empty dot) and GC-treated (filled dot) cells depleted of *PAXIP1* expression. Mean values ± SD are depicted. Left and right are two biological replicates with three technical replicates each. Two-way ANOVA test was performed. F. Western blot analyses showing GR expression, with actin as loading control, in NT, *PAXIP1*-KO, *STAG2*-KO, and *GR*-KO cells (n=2). Empty and filled dots indicate treatment with GCs for 24 hours.

**Figure Sup 2.**
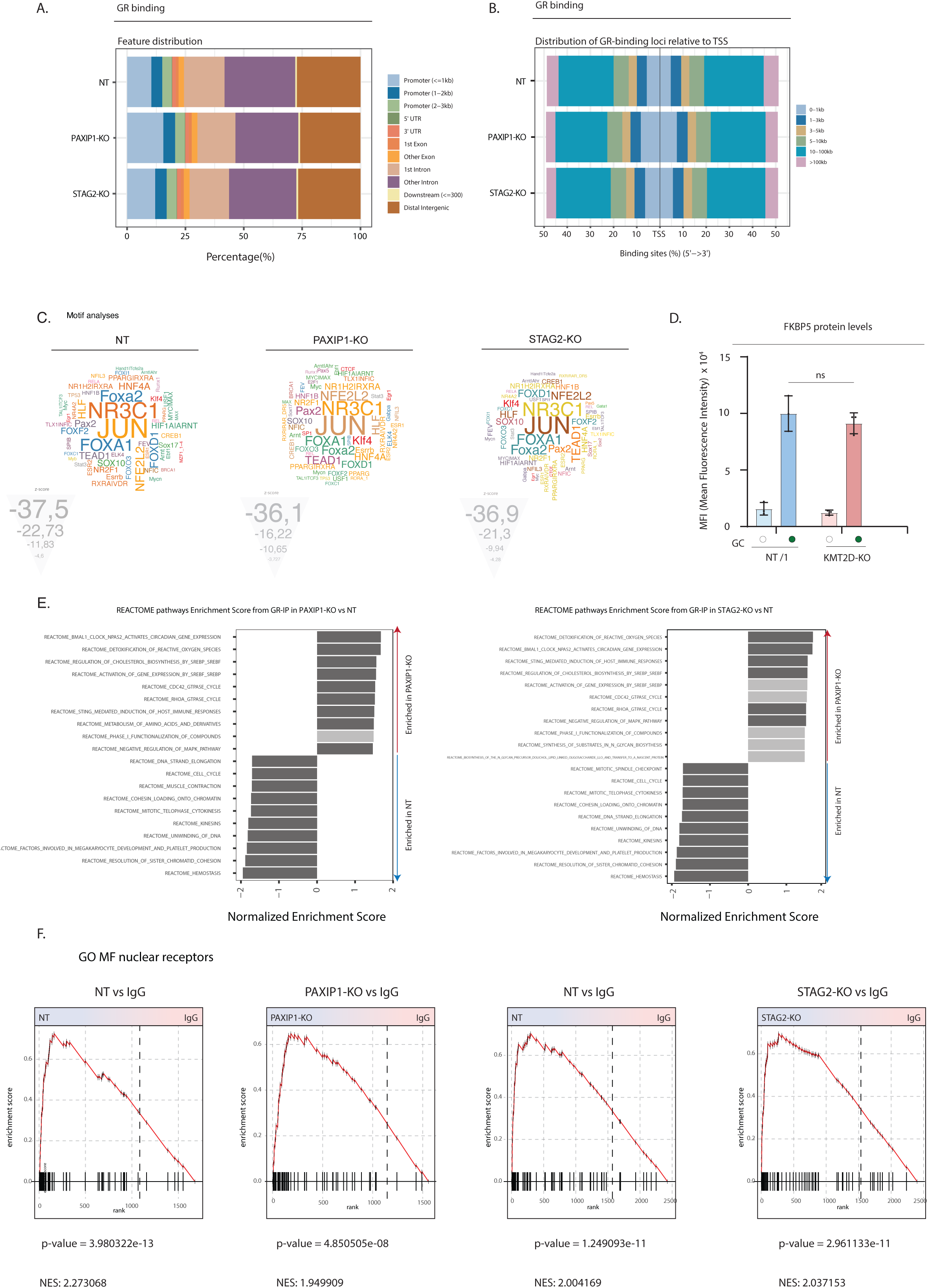
GR binding and GR complex composition in NT, *PAXIP1*-KO and *STAG2*-KO cells. A. Genomic distribution of GR binding in NT, *PAXIP1*-KO and *STAG2*-KO cells. B. Distribution of GR binding across the genome relative to the most-proximal transcription start site (TSS). C. Motif analyses of GR sites on NT (left), PAXIP1-KO (middle) and STAG2-KO (right) cells. Font size represents z-score. Motifs are colored according to different families of transcription factors. D. FKBP5 protein expression assessed by Flow Cytometry in NT and KMT2D-KOs. Empty and filled dots indicate treatment with DMSO or GCs for 24 hours, respectively. Quantification of Mean-Fluorescence Intensity (MFI) of FKBP5 expression (n=3). Two-way ANOVA test was performed. E. Gene Set Enrichment analyses of RIME data in *PAXIP1*-KO vs NT (left) and *STAG2*-KO vs NT (right), showing the top10 differentially expressed genesets. Dark grey coloring represents a nominal p-value < 0.05. F. Gene set enrichment profiles for nuclear transcription factor complex (GOMF_NUCLEAR_RECEPTOR_BINDING, GO:0016922) for NT-1, *PAXIP1*-KO, NT-2, *STAG2*-KO vs IgG. Nominal p-values and NES were calculated using GSEA.

**Figure Sup 3.**
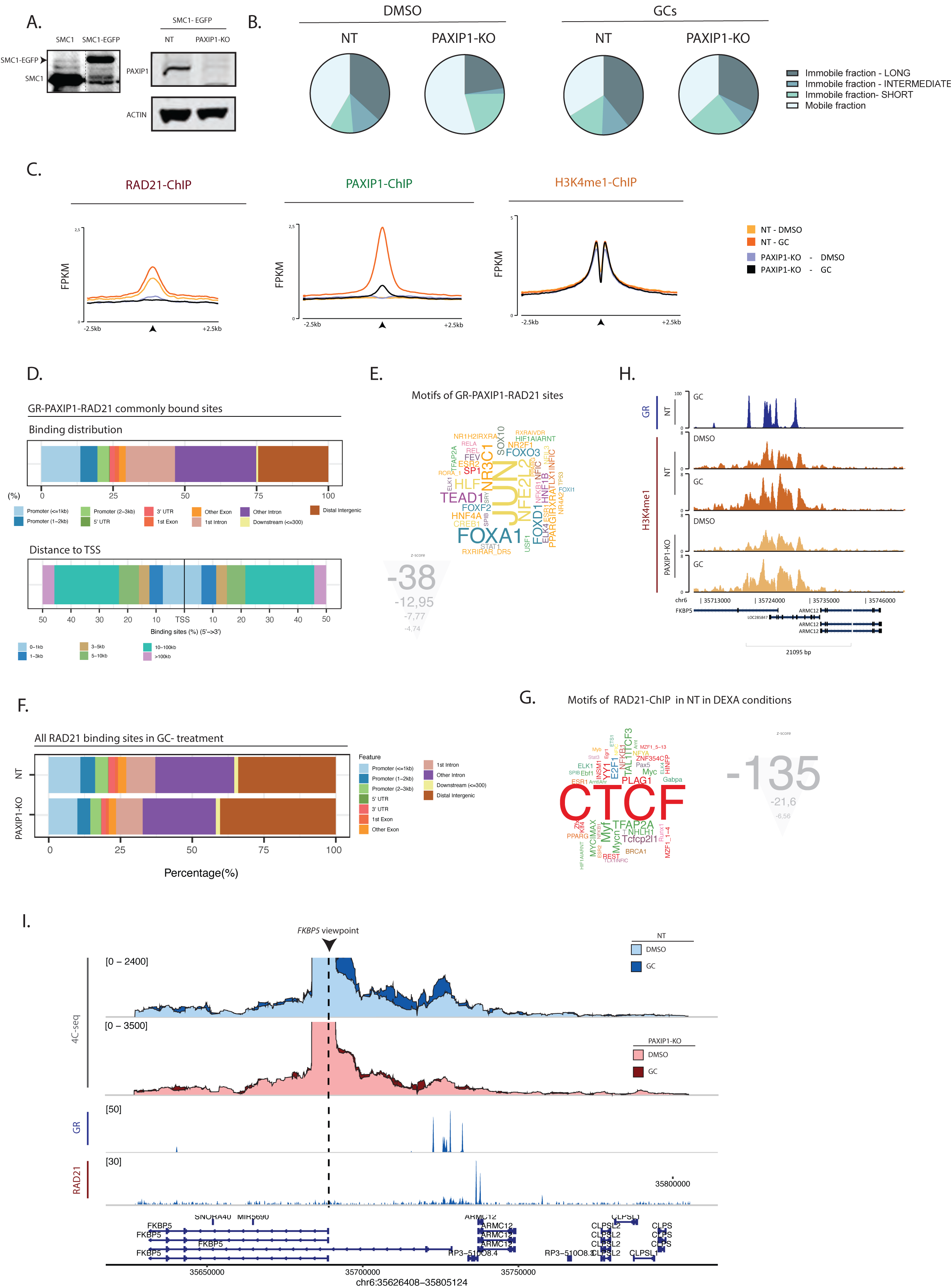
Studying the functional crosstalk between PAXIP1 and Cohesin. A. Western blot analyses showing SMC1-non-tagged and SMC1-EGFP tagged cells (left, n=2), and PAXIP1 expression in SMC1-EGFP tagged cells, with actin as control, in NT-SMC1-EGFP and *PAXIP1*-KO-SMC1-EGFP (n =2). B. Pie-charts representing calculated fraction sizes of mobile, and short, intermediate and long fractions attributed to SMC1-EGFP, in NT and PAXIP1-KO cells, in both DMSO and GC-treated cells. Monte Carlo simulation calculations were performed to calculate each fraction. C. Average density plot of RAD21 (left), PAXIP1 (middle) and H3K4me1 (right) ChIP-seq signal in NT and *PAXIP1*-KO cells in vehicle or GC-treated conditions, depicting a ±2.5kb window around the peak center. Data represent the average of two biological replicates. D. Genomic distribution (top) and distance to TSS (bottom) of GR-PAXIP1-RAD21 commonly bound sites. E. Motif analyses of GR-PAXIP1-RAD21 sites. Font size represents z-score. Motifs are colored according to different families of transcription factors. F. RAD21 genomic binding distribution in NT and *PAXIP1*-KO under GC-treated conditions. G. Motif analyses of RAD21-binding sites across all the genome. Font size represents z-score. Motifs are colored according to different families of transcription factors. H. ChIP-seq tracks at *FKBP5* locus depicting GR and H3K4me1 binding following GC treatment for 2 hours. Data is the merge of three (GR) and two (H3K4me1) independent biological replicates. A. 4C-seq experiments in NT and *PAXIP1*-KO cells using the *FKBP5* TSS as a viewpoint showing DMSO and GC-treated conditions. 4C data is shown as the mean of three independent biological experiments. ChIP tracks of RAD21 and GR in GC-treated cells are depicted. ChIP-tracks are a representative snapshot of one biological replicate.

**Figure Sup 4.**
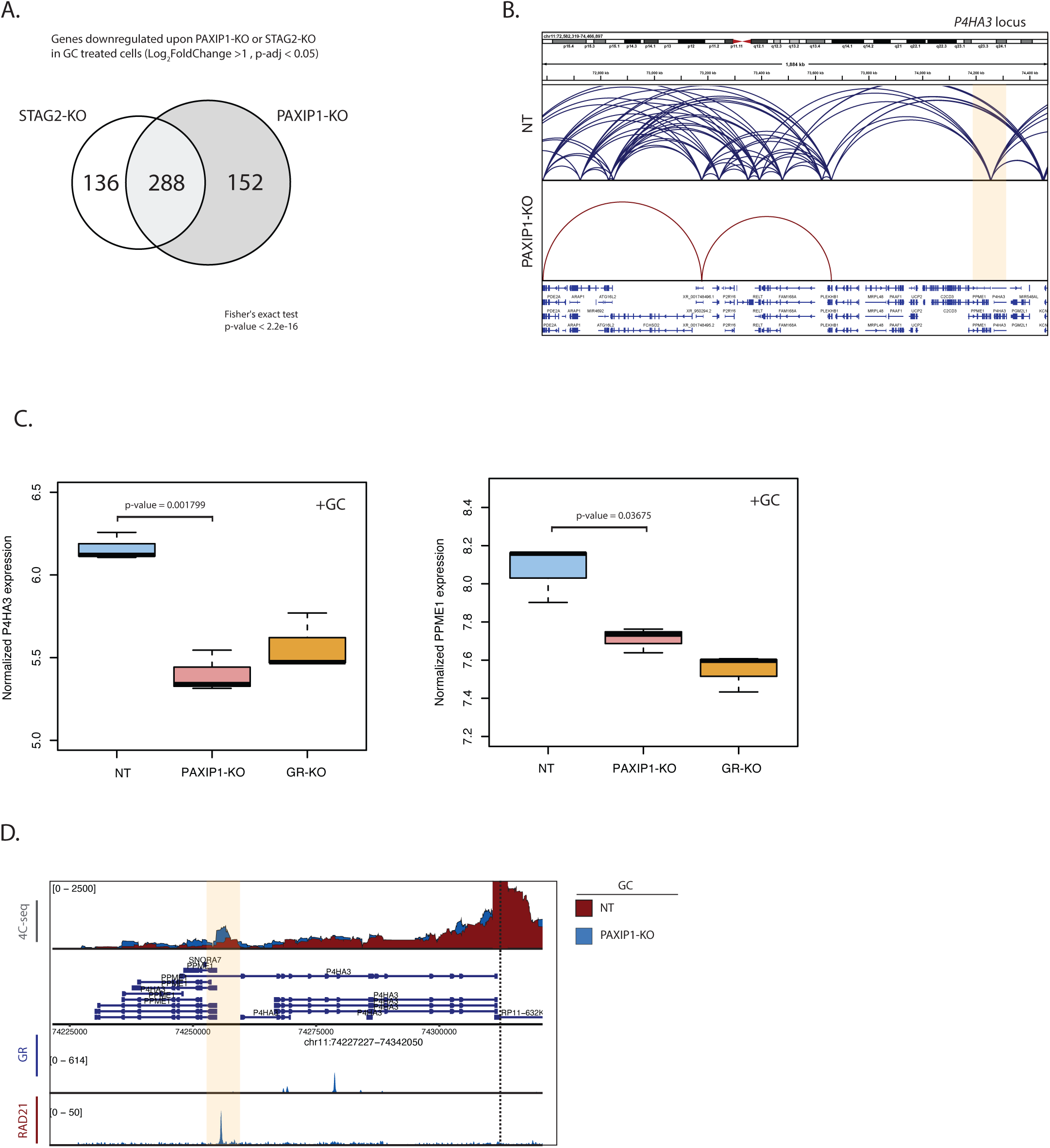
Expression of GR-target genes is dependent on enhancer-promoter interactions maintained by PAXIP1. A. Significantly downregulated genes with a log_2_(fold change)>1, and p-adjusted <0.05, in *PAXIP1*-KO and *STAG2*-KO cells treated with GC when compared to NT GC-treated cells. Significant overlap was calculated with a Fisheŕs exact t-test. B. Snapshot of sevenC prediction around the *P4HA3* and *PPME1* loci, both in NT and *PAXIP1*-KO cells following GC-treatment. C. Normalized *P4HA3* (left) and *PPME1* (right) expression in NT, *PAXIP1*-KO and *GR*-KO cells (n = 3). p-value was calculated using U Mann-Whitney t-test. D. 4C-seq experiments in NT and *PAXIP1*-KO cells using the *P4HA3* TSS as a viewpoint showing NT vs *PAXIP1*-KO in GC-treated conditions. 4C data is shown as the mean of three independent biological experiments. ChIP tracks of RAD21 and GR in GC-treated cells are depicted. ChIP-tracks are a representative snapshot of one biological replicate.

**Figure Sup 5.**
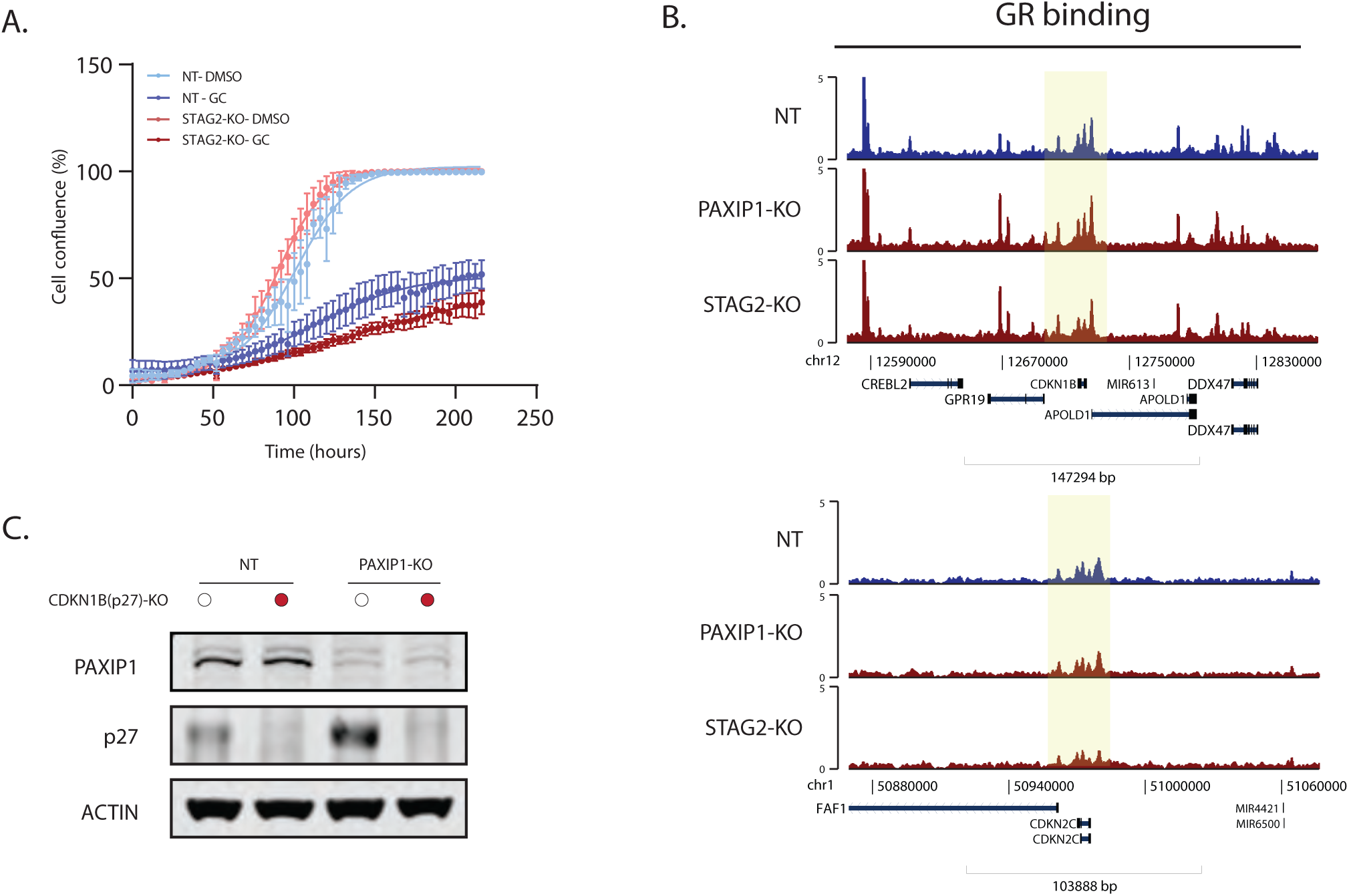
*PAXIP1* and *STAG2* loss enhances GC-sensitivity by increasing *CDKN1B* expression. A. Cell proliferation of NT and *STAG2*-KO cells following DMSO or GC treatment over time (n = 3 up to 168 hours, n = 2 from 168 hours to 216 hours, error bars represent 95% confidence interval). B. GR binding to *CDKN1B* and *CDKN2C* loci in NT, *PAXIP1*-KO, *STAG2*-KO cells in GC-treated conditions. Data are representative of three independent biological experiments. C. Western blot analyses showing PAXIP1 and P27 expression in NT, *PAXIP1*-KO and *PAXIP1*-KO/*CDKN1B*-KO DKO cells, with actin as a loading control (n =2). Actin has been used as loading control.

## Notes

### Competing Interest Statement

The authors have declared no competing interest.

